# Flow-Mediated Modulation of the Endothelial Cell Lipidome

**DOI:** 10.1101/2024.06.13.598934

**Authors:** Soon-Gook Hong, John P. Kennelly, Kevin J. Williams, Steve J. Bensinger, Julia J. Mack

## Abstract

The luminal surface of the endothelium is exposed to dynamic blood flow patterns that are known to affect endothelial cell phenotype. While many studies have documented the phenotypic changes by gene or protein expression, less is known about the role of blood flow pattern on the endothelial cell (EC) lipidome. In this study, shotgun lipidomics was conducted on human aortic ECs (HAECs) exposed to unidirectional laminar flow (UF), disturbed flow (DF), or static conditions for 48 hrs. A total of 520 individual lipid species from 17 lipid subclasses were detected. Total lipid abundance was significantly increased for HAECs exposed to DF compared to UF conditions. Despite the increase in the total lipid abundance, HAECs maintained equivalent composition of each lipid subclass (% of total lipid) under both DF and UF. However, by lipid composition (% of total subclass), 28 lipid species were significantly altered between DF and UF. Complimentary RNA sequencing of HAECs exposed to UF or DF revealed changes in transcripts involved in lipid metabolism. Shotgun lipidomics was also performed on HAECs exposed to pro-inflammatory agonists lipopolysaccharide (LPS) or Pam3CSK4 (Pam3) for 48 hrs. Exposure to LPS or Pam3 reshaped the EC lipidome in both unique and overlapping ways. In conclusion, exposure to flow alters the EC lipidome and ECs undergo stimulus-specific lipid reprogramming in response to pro-inflammatory agonist exposure. Ultimately, this work provides a resource to profile the transcriptional and lipidomic changes that occur in response to applied flow that can be accessed by the vascular biology community to further dissect and extend our understanding of endothelial lipid biology.

## INTRODUCTION

Endothelial cells (ECs) line the luminal surface of the vasculature and are constantly exposed to mechanical forces generated by blood flow ^1^. Due to variations in vessel geometry and curvature, the arterial endothelium experiences different blood flow patterns, including unidirectional laminar flow (UF) generated in linear regions of the vasculature and oscillatory disturbed flow (DF) in regions of the vasculature that are highly curved or bifurcated ^2^. Different levels of hemodynamic shear stress (the frictional force acting on ECs) are present and range from high arterial shear stress (>15 dyne/cm^2^) induced by UF to low time-averaged shear stress (4 dyne/cm^2^) and oscillating flow direction induced by DF.

These distinct blood flow patterns are known to induce different EC phenotypes. While UF promotes a state of endothelial quiescence and an anti-inflammatory phenotype, a pro-inflammatory phenotype is associated with ECs exposed to DF ^3^. Therefore, exposure to DF presents as a local risk factor for vascular inflammation and disease progression ^4,5^. Furthermore, the endothelium exposed to DF is more susceptible to the development of chronic inflammatory diseases such as atherosclerosis ^2,4^.

Lipids are a major class of biological molecules that are essential for cellular function. Lipids are involved in many bioenergetic, biochemical, and biophysical processes ^6^ and so maintaining lipid homeostasis is critical to cellular health. All cellular membranes are composed of lipids, predominantly phospholipids including glycerophospholipids and sphingolipids ^7,8^. Neutral lipids, including triglycerides (TG) and cholesteryl esters, are also stored in cytosolic lipid droplets surrounded by a monolayer of phospholipids ^9^. In ECs, it has been demonstrated previously that mechanical forces due to applied flow affect the membrane lipid order and fluidity ^10^ and alter membrane cholesterol levels ^11^. However, a comprehensive assessment of the changes that occur to the EC lipidome in response to mechanical stress is lacking. In the last decades, many studies have set out to understand the mechanisms of EC inflammatory activation and to identify molecular pathways and potential therapeutic targets to alleviate vascular inflammation ^12^. However, the role of membrane lipids in the context of EC signaling and vascular inflammation has been less studied.

Here, we conducted complementary studies to characterize the transcriptional and lipidomic profiles of human aortic ECs (HAECs) exposed to the physiological flow conditions of UF and DF. We further compared these datasets to the lipidomic signatures to HAECs exposed to pro-inflammatory agonists. Our results provide a resource to examine how blood flow pattern affects the EC lipid profile and future work will hopefully lead to the identification of lipid species involved in endothelial inflammation and vascular disease.

## METHODS

### Cell culture and flow application

Primary HAECs (Cell Applications #S304-05a, Lot#1487(s); healthy normal human aorta from 21-year-old Caucasian male) were used from P4 to P12. For cell culture experiments, MCDB-131 complete media (VEC Technologies #MCDB-131 Complete) was supplemented with 10% FBS (Omega USDA certified FBS #FB-11). For plating cells on cell culture dishes, 0.1% gelatin (Stemcell #07903) coating was first applied. Cells were cultured in a 37 °C incubator with 5% CO_2_. For application of shear stress, HAECs were seeded in ibidi µ-Slide 0.2 (ibidi #80166) or 0.6 (ibidi #80186) Luer ibiTreat. Unidirectional laminar flow (20 dynes/cm^2^) or disturbed flow (±4 dynes/cm^2^, 2 Hz) was applied to confluent monolayers using the ibidi pump system (ibidi #10902) for 48 hrs.

### RNA sequencing analysis

HAECs were seeded on ibidi µ-Slides and subjected to unidirectional laminar flow (20 dynes/cm^2^) or disturbed flow (±4 dynes/cm^2^, 2 Hz) using the ibidi pump system for 48 h in a 37°C incubator with 5% CO_2_. After 48 h, cells were washed with 1X PBS (Gibco #14190-144) and then lysed using RLT lysis buffer (QIAGEN #79216) plus 1% 2-Mercaptoethanol (Sigma Aldrich #M6250). For each condition, four µ-Slides were combined as one flow-exposed sample for a total of 12 slides for n = 3 samples. RNA extraction, library construction, and sequencing were conducted by the UCLA Technology Center for Genomics & Bioinformatics (TCGB) core facility.

Briefly, libraries for RNA-Seq were prepared with KAPA Stranded mRNA-Seq Kit (KAPA Biosystems #KK8421). Briefly, the workflow consisted of mRNA enrichment and fragmentation, first strand cDNA synthesis using random priming followed by second strand synthesis converting cDNA:RNA hybrid to double-stranded cDNA (ds-cDNA), and incorporation of dUTP into the second cDNA strand. cDNA generation was followed by end repair to generate blunt ends, A-tailing, adaptor ligation and PCR amplification. Different adaptors were used for multiplexing samples in one lane. Sequencing was performed on Illumina HiSeq 3000 for SE 1x65bp run. Data quality check was done on Illumina SAV. Demultiplexing was performed with Illumina Bcl2fastq v2.19.1.403 software. The alignment was performed using Spliced Transcripts Alignment to a Reference (STAR) ^13^ using the human reference genome assembly GRCh38. The Ensembl Transcripts release GRCh38.107 GTF was used for gene feature annotation. For normalization of transcripts counts, counts per million (CPM) normalized counts were generated by adding 1.0E-4 followed by by counts per million (CPM) normalization.

For graphical display of differential expression, Heatmapper ^14^ was used to create a heatmap of gene expression. Average linkage was used as Clustering Method, and Kendall’s Tau was used as Distance Measurement Method. VolcaNoseR was used to generate the Volcano plot of gene expression. Log_2_ (fold change; FC) as X-variables and –log10(p-value) as Y-variables were used. ShinyGO 0.77 ^15^ was used for gene enrichment analysis with false discovery rate (FDR) of 0.05 as cut-off. The Database for Annotation, Visualization and Integrated Discovery (DAVID, NIH) was used to identify the biological pathways enriched under DF vs. UF. Principal component analysis (PCA) was conducted using ClustVis ^16^.

Gene Set Enrichment Analysis (GSEA) 4.3.2 software was also used to identify biological pathways enriched in response to DF. Genes used for GSEA were those that were differentially expressed in HAECs under DF versus UF. The Hallmarks gene set database was used with the following analysis parameters: Number of permutations: 1000; Collapse/Remap to Gene Symbols: No Collapse; Permutation type: Gene Set; Chip Platform: Human Gene Symbol with Remapping MSigDB.v2023.1.Hs.chip; Enrichment Statistic: Weighted; Metric for Ranking Genes: Signal2Noise; Gene List Sorting Mode: Real; Gene List Ordering Mode: Descending; Max Size: Exclude Larger Sets: 500; Min Size: Exclude Smaller Sets: 15.

From the Gene Ontology (GO) database, lists of the genes involved in lipid catabolic process (GO:0016042), neutral lipid catabolic process (GO:004646), and fatty acid catabolic process (GO:0009062) were identified from using the RNAseq dataset.

### Lipidomics analysis

For flow experiments, HAECs were seeded on ibidi µ-Slides and subjected to unidirectional laminar flow (20 dynes/cm^2^) or disturbed flow (±4 dynes/cm^2^, 2 Hz) using the ibidi pump system for 48 h in a 37°C incubator with 5% CO_2_. After 48 h, cells were washed with ice-cold 1X PBS and subjected to trypsinization for cell detachment from the slide. For each flow condition, 3 µ-Slides were combined as one sample for a total of 9 slides for n = 3 samples. For inflammatory agonist treatments, HAECs were seeded on 6-well plates (Corning #353046) and treated with either LPS (50ng/ml; Invivogen, #tlrl-smlps), Pam3CSK4 (200 ng/ml; Invivogen #tlrl-pms), or vehicle control for 48 hrs. Cells were collected and resuspended after centrifugation at 900 rpm for 5 min at 4°C. Cell number was assessed by hemocytometer. Lipid extraction and analysis were conducted by the UCLA Lipidomics Lab.

Cells were transferred to extraction tubes with 1X PBS. A modified Bligh and Dyer extraction ^17^ was carried out on samples. Prior to biphasic extraction, a standard mixture of 75 lipid standards (Avanti #330820, 861809, 330729, 330727, 791642) was added to each sample. Following two successive extractions, pooled organic layers are dried down in a Thermo SpeedVac SPD300DDA using ramp setting 4 at 35°C for 45 minutes with a total run time of 90 min. Lipid samples were resuspended in 1:1 methanol/dichloromethane with 10mM ammonium acetate and transferred to robovials (Thermo #10800107) for analysis.

Samples were analyzed by direct infusion on a Sciex 5500 with Differential Mobility Device (DMS) — comparable to the Sciex Lipidyzer platform—with a targeted acquisition list consisting of 1450 lipid species across 17 subclasses. The DMS was tuned with EquiSPLASH LIPIDOMIX (Avanti #330731). Data analysis was performed with in-house data analysis workflow. All instrument settings, MRM lists, and analysis methods are available ^18^. Quantitative values were normalized to cell counts.

Heatmaps were generated using Heatmapper ^14^ and ClustVis ^16^. Principal component analysis (PCA) was conducted using ClustVis. X and Y axis show principal component 1 (PC1) and principal component 2 (PC2), respectively. PCA loadings were extracted from the ClustVis, and top 20 loadings for PC1 were identified. Cross analysis of the datasets was conducted using InteractiVenn ^19^ to identify lipid species that were changed under both DF and pro-inflammatory agonist exposure.

### Gene expression analysis

For gene expression analysis using qPCR, using RNeasy Mini kit (QIAGEN #74104), total RNA was extracted from HAECs. RNA was converted to cDNA by reverse transcription, and cDNA was quantified using a Nano-Drop 8000 (Thermo Fisher). Target genes were quantified using iTaq Universal SYBR Green Supermix (BioRad #1725125) and the appropriate primer pairs. Samples were run on a QuantStudio 6 Flex 384-well qPCR apparatus (Applied Biosystems). Each target gene was normalized to HPRT. Primer sequences (5′ – 3′) are listed below.

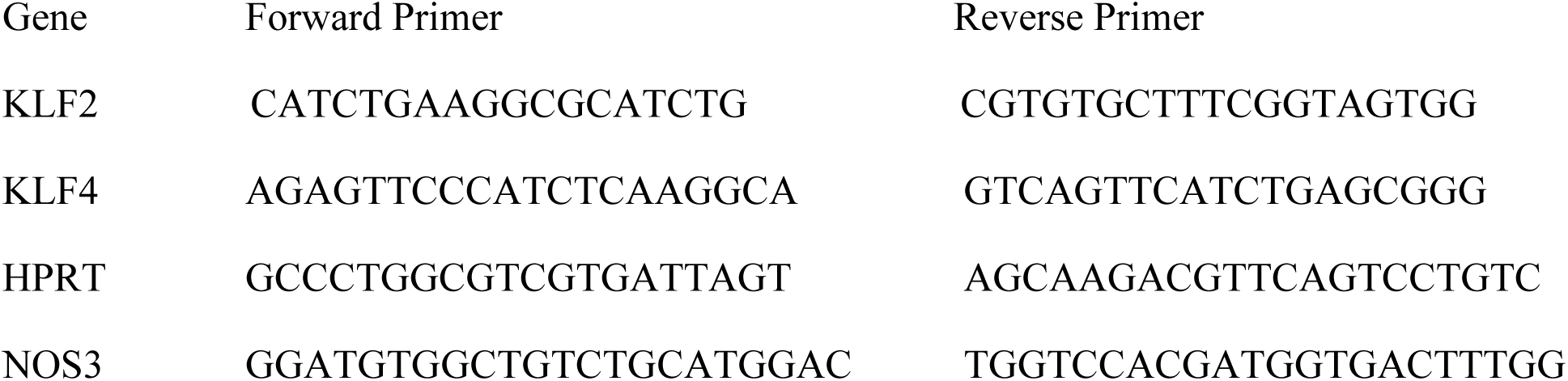

### Immunostaining and confocal imaging

For immunostaining, HAECs were fixed with 4% PFA for 5 min followed by multiple washes with 1X PBS. Samples were then blocked for 1 h with 5% Normal donkey serum (Jackson Immuno Research Laboratories #017-000-121) in 1X PBS. Then, samples were permeabilized with 0.1% Triton X-100 (Fisher Scientific #A16046-0F) for 5 min followed by blocking with 5% Normal donkey serum for 1 hr. Primary antibodies were incubated overnight at 4 °C in blocking buffer and secondary antibodies applied for 1 h at room temperature. Primary antibodies used: NOS3 (Santa Cruz #sc-376751), ICAM-1 (Santa Cruz #sc-107), NF-κB p65 (Cell Signaling #8242S), VE-Cadherin (R&D #AF938). Imaging was performed on a Zeiss LSM 900 confocal microscope equipped with 405nm, 488nm, 561nm and 640nm laser lines using Plan-Apochromat objectives (20x) and Airyscan2 GaAsP-PMT detector. Identical laser intensity settings were applied to all samples being compared with equivalent Z thickness. After acquisition, a maximum intensity projection of the Z-stack was applied using ZEN Blue 3.5 software (ZEISS). Image processing and quantification of parameters was performed with ImageJ (NIH).

### Drawings

Schematics in figures were created using BioRender.com.

### Statistical analysis

Statistical analysis was performed using GraphPad Prism software. The results are presented as mean ± SD. The Shapiro-Wilk normality test was conducted to measure normal distribution of dependent variables. For normally distributed data, two-tailed independent t-test was used to determine statistically significant differences between two groups. For data that did not pass the normality test, nonparametric one-tailed Mann-Whitney U test was conducted to examine statistical significance between 2 experimental conditions. p< 0.05 was considered statistically significant for all analyses.

## RESULTS

### Disturbed flow alters genes associated with lipid metabolism

HAECs were exposed to UF (20 dynes/cm^2^) or DF (±4 dynes/cm^2^, 2 Hz) for 48 hrs and subjected to RNA sequencing (RNAseq) and shotgun lipidomics **(Figure 1A)**. We confirmed the expected difference in cell morphology of HAECs after exposure to UF or DF for 48 h **(Figure 1B).** To confirm that our in vitro flow system generated the expected EC phenotypes under the applied flow conditions, we validated both gene and protein expression. Gene expression of known flow-responsive genes, including KLF2, KLF4, and NOS3 showed a significant upregulation under UF **(Supplemental Figure 1A)**. And confocal imaging of HAEC monolayers confirmed the increase in eNOS protein expression levels under UF **(Supplemental Figure 1B)**.

**FIGURE 1:**
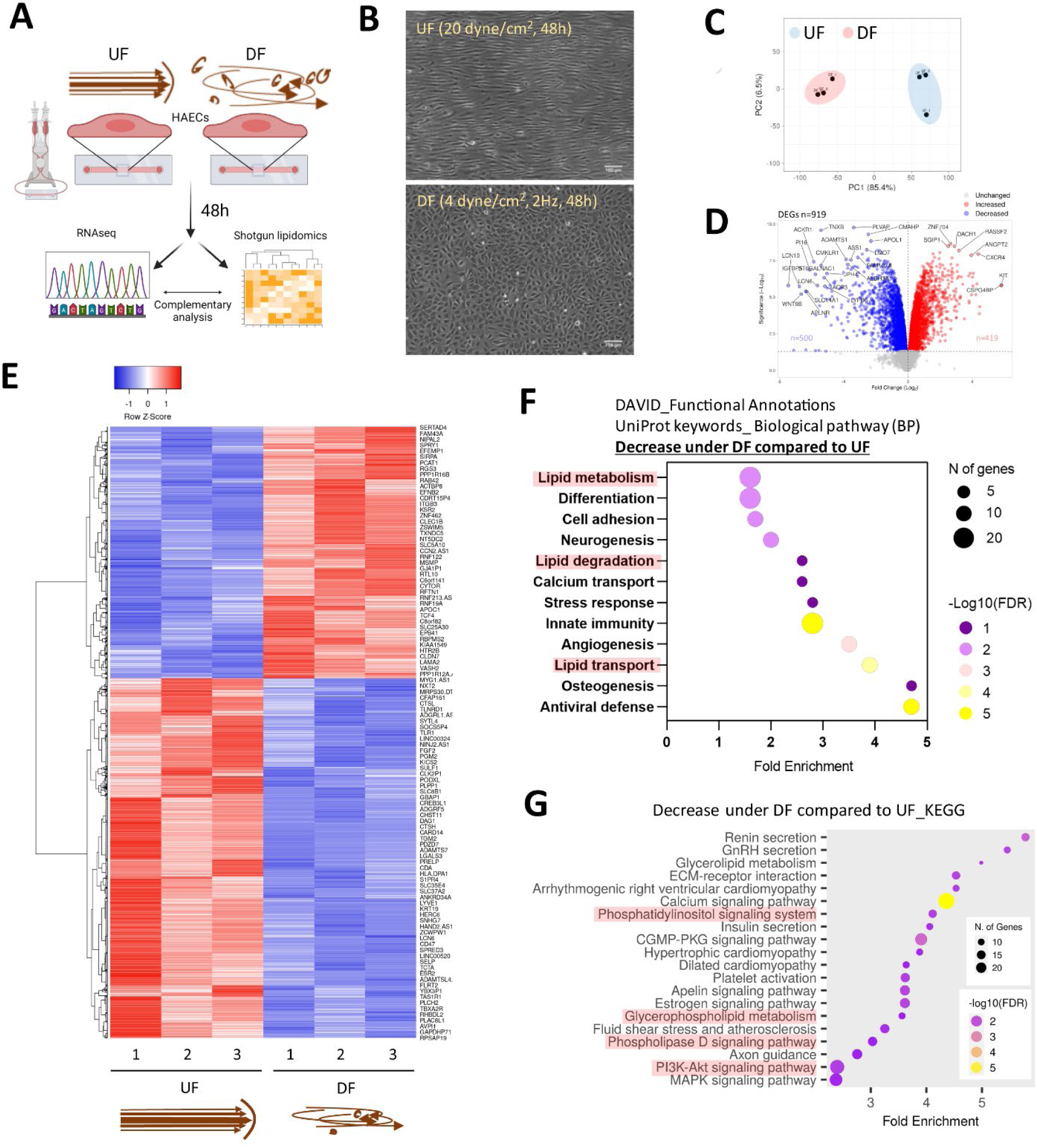
Flow pattern affects the transcriptional signature of HAECs and alters the expression of genes related to lipid metabolism. **(A)** Schematic diagram of the experimental procedure. (B) Brightfield micrographs of HAEC monolayers exposed to UF and DF for 48 h. **(C)** PCA plot of RNA sequencing (RNASeq) data. Blue dots = UF-exposed HAECs. Red dots = DF-exposed HAECs. **(D)** Volcano plot of differentially expressed genes (DEGs). **(E)** Heatmap of DEGs. **(F)** Pathway analysis using DAVID (UniProt database) showing the biological processes downregulated under DF vs. UF. **(G)** Pathway analysis using ShinyGO (KEGG database) to show reduced biological pathways under DF vs. UF. UF = Unidirectional laminar flow. DF = Disturbed flow.

Consistent with previous reports ^20^, our RNAseq analysis showed that HAECs exposed to DF resulted in dramatic changes in the gene expression profile compared to HAECs exposed to UF **(Figure 1 C–E)**. Pro-inflammatory genes known to be induced by DF, such as Angiopoietin 2 (*ANGPT2*), Endothelin 1 (*EDN1*), C-C Motif Chemokine Ligand 2 (*CCL2*), NADPH Oxidase 4 (*NOX4*), 6-phosphofructo-2-kinase/fructose-2,6-biphosphatase 3 (*PFKFB3*), and Hypoxia-Inducible-Factor 1A (*HIF1A*) ^21–23^, were significantly elevated in HAECs exposed to DF **(Supplemental Figure 1C)**. Additionally, we confirmed the pro-inflammatory phenotype of HAECs exposed to DF by confocal imaging. HAECs exposed to DF had increased nuclear NF-κB p65 localization and ICAM-1 protein expression compared to HAECs exposed to UF **(Supplemental Figure 1D & 1E)**. Pathway analysis of differentially expressed genes for HAECs exposed to DF vs. UF revealed changes in transcripts associated with lipid metabolism, lipid degradation, lipid transport, and lipid signaling (**Figure 1F & 1G**).

### Exposure to disturbed flow increases total cellular lipid content

Based on the observation that genes associated with lipid metabolism were altered in DF compared to UF, we hypothesized that the applied flow pattern may affect the EC lipidome. Therefore, we compared the lipid profiles of HAECs exposed to UF and DF for 48 hrs **(Figure 2A)**.

**FIGURE 2:**
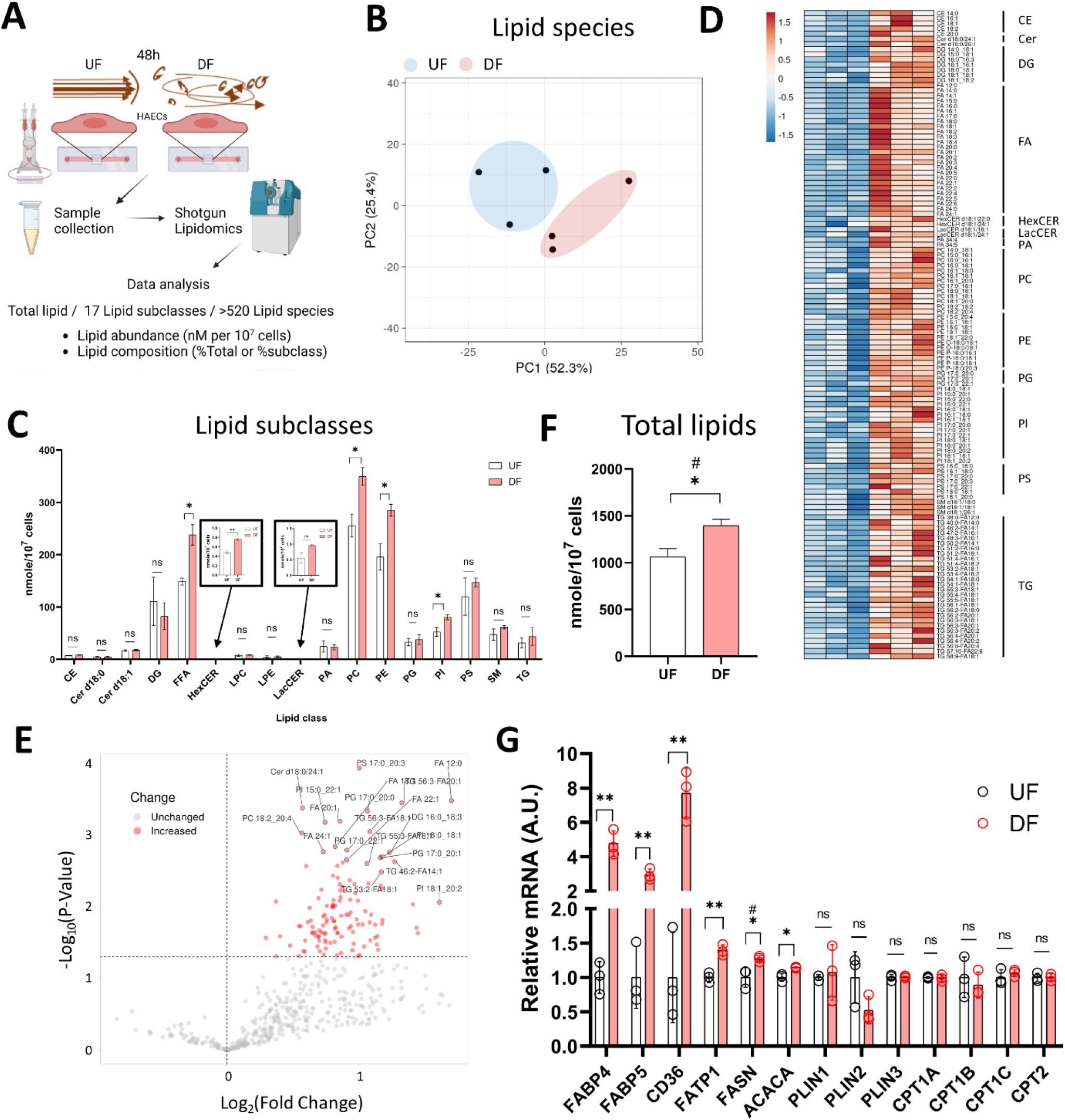
Lipid abundance is elevated in HAECs exposed to DF compared to UF. **(A)** Schematic diagram of the lipidomics analysis. **(B)** PCA plot of lipidomics data. Blue dots = UF-exposed HAECs. Red dots = DF-exposed HAECs. **(C)** Abundance of each lipid subclasses under UF vs. DF (n=3). **(D)** Heatmap of altered lipid species for DF vs. UF. **(E)** Volcano plot showing altered lipid species under DF vs. UF. **(F)** Total lipid abundance (nM per 10^7^ cells) under UF vs. DF. Total lipid abundance was calculated from lipidomics results (n=3). **(G)** Expression of the genes involved in fatty acid synthesis, uptakeand transport from the RNAseq data (n=3). Bar graph shown as mean ± SD. *p<0.05, **p<0.01, ns=not significant by two-tailed unpaired *t*-test. #p<0.05 by one-tailed Mann-Whitney U test.

Shotgun lipidomics detected a total of 520 individual lipid species from 17 lipid subclasses **(Figure 2A)**. As visualized by principal component analysis (PCA), the lipid profile was distinct between HAECs exposed to UF and DF **(Figure 2B)**. Of the 17 lipid subclasses measured, 5 lipid subclasses were significantly elevated in HAECs exposed to DF compared to UF (**Figure 2C**). The lipid subclasses that were higher in DF compared to UF included free fatty acids (FFA), hexosylceramides (HexCER), phosphatidylcholines (PC), phosphatidylethanolamines (PE), and phosphatidylinositols (PI). PC and PE were the most abundant lipid subclasses detected for HAECs under both flow profiles (**Figure 2C**). A comparison of individual lipid species showed that 66 species were significantly elevated under DF versus UF **(Figure 2D & 2E)**. Furthermore, the total lipid abundance per cell was higher for HAECs exposed to DF compared to UF (**Figure 2F**).

Further analysis indicated an increase in the abundance of unesterified fatty acids for HAECs exposed to DF **(Supplemental Figure 2A & 2B)**. Fatty acyl chains for phospholipid and glycerolipid synthesis are obtained via import or endogenous synthesis ^24^ and the fatty acid abundance in cells is affected by proteins that regulate the uptake, synthesis, and transport of fatty acids, as well as proteins that hydrolyze fatty acids from neutral lipids and phospholipids (**Supplemental Figure 2D**). Cross analysis of the RNAseq data indicated that genes involved in fatty acid synthesis (*ACACA*, *FASN*), fatty acid uptake (*CD36*) and fatty acid transport (*FABP4*, *FABP5*, *FATP1)* were upregulated under DF (**Figure 2G**). Interestingly, genes involved in lipid catabolic processes, neutral lipid catabolic processes, and fatty acid catabolic processes tended to be reduced in the presence of DF **(Supplemental Figure 3 A–D)**. Additionally, GSEA analysis revealed an increase in transcripts related to cholesterol biosynthesis under DF **(Supplemental Figure 3E)**, further suggesting that the type of flow modulates lipid metabolic pathways in HAECs. Together, gene expression analysis indicated a shift towards a program of increased lipid biosynthesis and import with reduced lipid turnover.

Comparing HAECs under static conditions revealed a lipid profile quite distinct from HAECs exposed to either DF or UF **(Supplemental Figures 4 & 5)**. PCA of the lipidomics data showed that cells exposed to UF or DF were clearly separated from cells in static conditions on PC1, and cells exposed to DF or static conditions were separated from cells exposed to UF on PC2 **(Supplemental Figure 6A)**. The top 20 lipid species responsible for the separation along PC1 and PC2 are listed in **Supplemental Figure 6B**. Furthermore, HAECs exposed to DF had more lipid content per cell than HAECs exposed to either UF or static conditions (**Supplemental Figure 6C**). A heatmap of differentially expressed lipid species showed clear effects of applied flow on the lipid profile of HAECs **(Supplemental Figure 6D)**.

### Specific lipid species are altered by disturbed flow

We next analyzed the lipid composition of cells exposed to DF or UF by expressing each lipid subclass as a percent (%) of total cellular lipids. Despite the increase in total lipid mass per cell in the DF condition, the percent composition of each lipid subclass remained unchanged for HAECs exposed to DF or UF (e.g., PCs accounted for ~25% total cellular lipids in both DF and UF) **(Figure 3A)**. Therefore, we examined whether there was a shift in the contribution of specific lipid species as a fraction of its total subclass. Twenty-three lipid species were increased, and five lipid species were decreased under DF compared to UF **(Figure 3B–E).** A closer look at the lipid species indicated that phospholipids containing mono-unsaturated fatty acids (MUFA) at the sn-2 position were increased in DF compared to UF (e.g., PE 18:1_18:1, PG 16:0_18:1, PG 18:1_18:1, PI 18:0_18:1, PI 18:0_20:1). Conversely, phospholipid species containing polyunsaturated fatty acids (e.g., PC 18:0_20:4, PE 18:0_20:4, PE 18:0_22:4) were decreased under DF.

**FIGURE 3:**
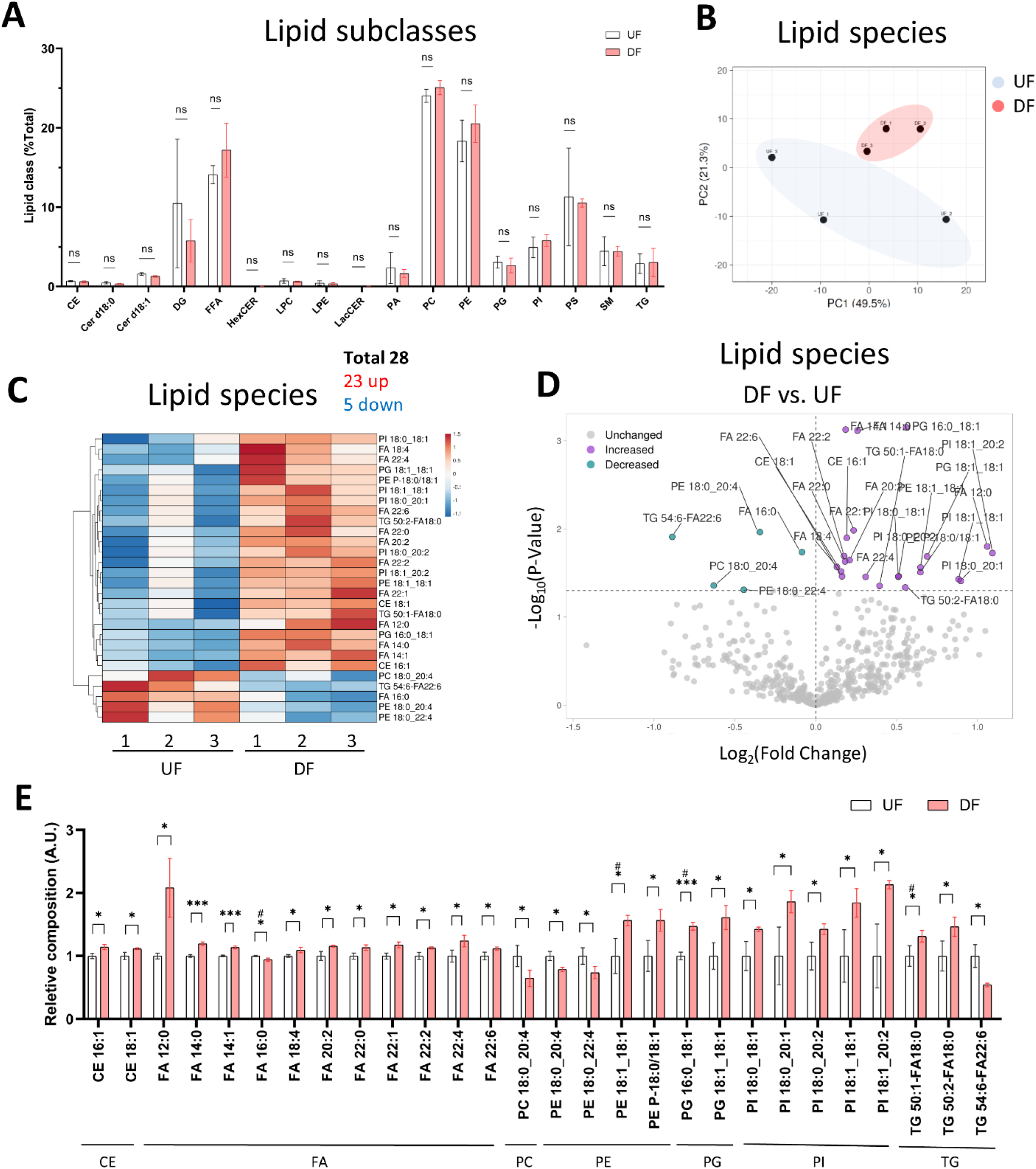
Lipid subclass composition is not changed by flow pattern. **(A)** Bar graph of lipid subclass composition (lipid subclass / total lipid) for HAECs exposed to UF and DF (n=3). **(B)** PCA plot using lipid species composition. **(C)** Heatmap of lipid species changed by percent composition under DF vs. UF. **(D)** Volcano plot of lipid species altered in composition. **(E)** Bar graph of lipid species altered in composition under DF vs. UF (n=3). Bar graph shown as mean ± SD. *p<0.05, **p<0.01, ***p<0.001, ns=not significant by two-tailed unpaired *t*-test. #p<0.05 by one-tailed Mann-Whitney U test.

### Endothelial lipid profile is altered in response to pro-inflammatory signals

Since DF results in an inflammatory EC phenotype (**Supplemental Figure 1 C–E**), we asked if specific pro-inflammatory signals also altered the lipid composition of ECs. To address this, we performed shotgun lipidomics on statically cultured HAECs in the absence or presence of pro-inflammatory agonists Pam3CSK4 (Pam3; Toll-like receptor 1/2 agonist) or lipopolysaccharides (LPS; Toll-like receptor 4 agonist) for 48 hr. We first confirmed that exposure to LPS and Pam3 for 48 hr at the dosages applied resulted in an inflammatory phenotype by staining for ICAM-1 and NF-κB p65. As expected, HAECs exposed to LPS or Pam3 had increased ICAM-1 and nuclear localization of NF-κB p65 compared to control HAECs (**Supplemental Figure 7 A–D**). And as we observed for HAECs exposed to DF, the total lipid abundance per cell was significantly elevated in response to LPS and Pam3 compared to vehicle treated control cells **(Figure 4A)**. Cellular lipidomes were distinct between HAECs treated with pro-inflammatory agonists versus control HAECs **(Figure 4B, 4C, 4E, 4F)**. In summary, a total of twelve lipid subclasses were elevated in response to LPS and eight lipid subclasses were elevated in response to Pam3 (**Figure 4D**). We noted that cholesteryl ester (CE) and sphingomyelin (SM) subclasses were reduced in Pam3-treated HAECs, while CE and SM were increased in response to LPS **(Figure 4D)**. In addition, PCA and hierarchical clustering showed that exposure to inflammatory agonists led to dramatic differences in the cellular lipid composition **(Supplemental Figure 7 E-I)**.

**FIGURE 4:**
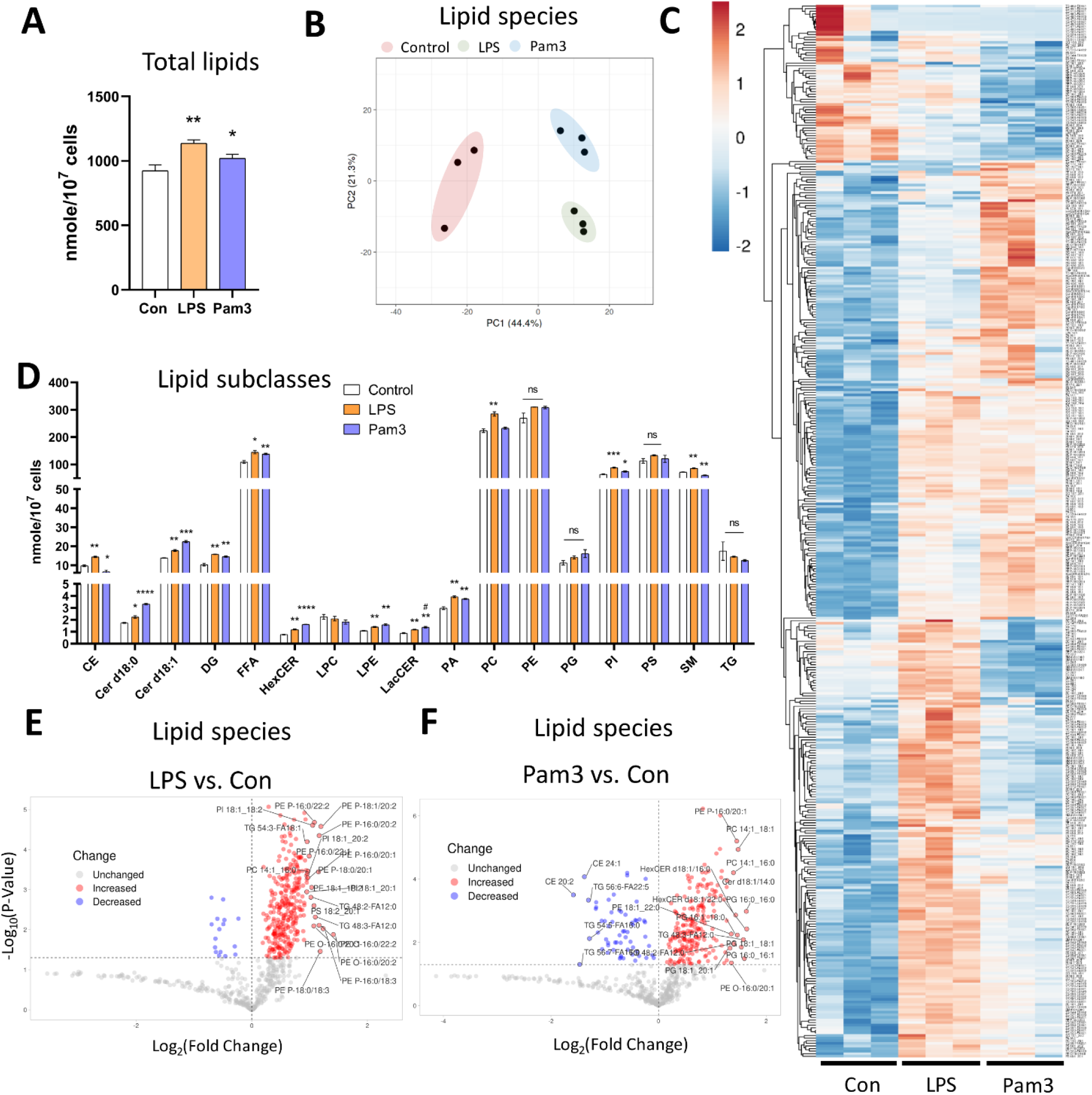
Lipid profile is altered in response to inflammatory agonist exposure. **(A)** Total lipid abundance (nM per 10^7^ cells) in HAECs treated with vehicle (control), LPS, and Pam3. Total lipid abundance was calculated from lipidomics results (n=3). (**B**) PCA plot. Red = control HAECs, Green = LPS-treated HAECs, Blue = Pam3-treated HAECs. **(C)** Heatmap of altered lipid species in composition in response to vehicle, LPS, and Pam3. **(D)** Bar plot of lipid subclass abundance in control, LPS, and Pam3 groups (n=3). **(E)** Volcano plot of lipid species changed in LPS vs. control. **(F)** Volcano plot of lipid species changed in Pam3 vs. control. Bar graph shown as mean ± SD. *p<0.05, **p<0.01, ***p<0.001, ns=not significant by two-tailed unpaired *t*-test. #p<0.05 by one-tailed Mann-Whitney U test.

### Specific lipid species are elevated in response to disturbed flow and inflammatory agonists

Cross-analysis of the datasets identified lipid species that were altered in the presence of both DF and inflammatory agonist exposure. The abundance of 57 lipid species (nMols per cell) were increased in either DF, LPS, or Pam3 exposure (**Supplemental Figure 8A**). However, by percent composition of individual lipid species, quantified as the percent abundance relative to the lipid subclass abundance, only six lipid species were increased in response to DF, LPS and Pam3 (**Supplemental Figure 8B**). Further analysis revealed that only five lipid species, specifically PE 18:1_18:1, PI 18:0_18:1, PI 18:0_20:2, PI 18:1_18:1, PI 18:1_20:2, were elevated in both abundance and percent composition in the presence of DF, LPS, and Pam3 **(Figure 5A**, **Figure 5C–F)**. We noted that stearic acid (18:0), oleic acid (18:1), and eicosadienoic acid (20:2), were common to these five lipid species. The total abundance of stearic acid, oleic acid, and eicosadienoic acid was elevated both in response to DF and inflammatory agonists **(Figure 5B)**. We further noted that four out of the five lipids that increased in absolute mass and percent of total lipid class by DF, LPS, and Pam3 were PI species. This observation was consistent with pathway enrichment scores for PI signaling and metabolism from the RNASeq data for HAECs exposed to DF **(Supplemental Figure 9)**.

**FIGURE 5:**
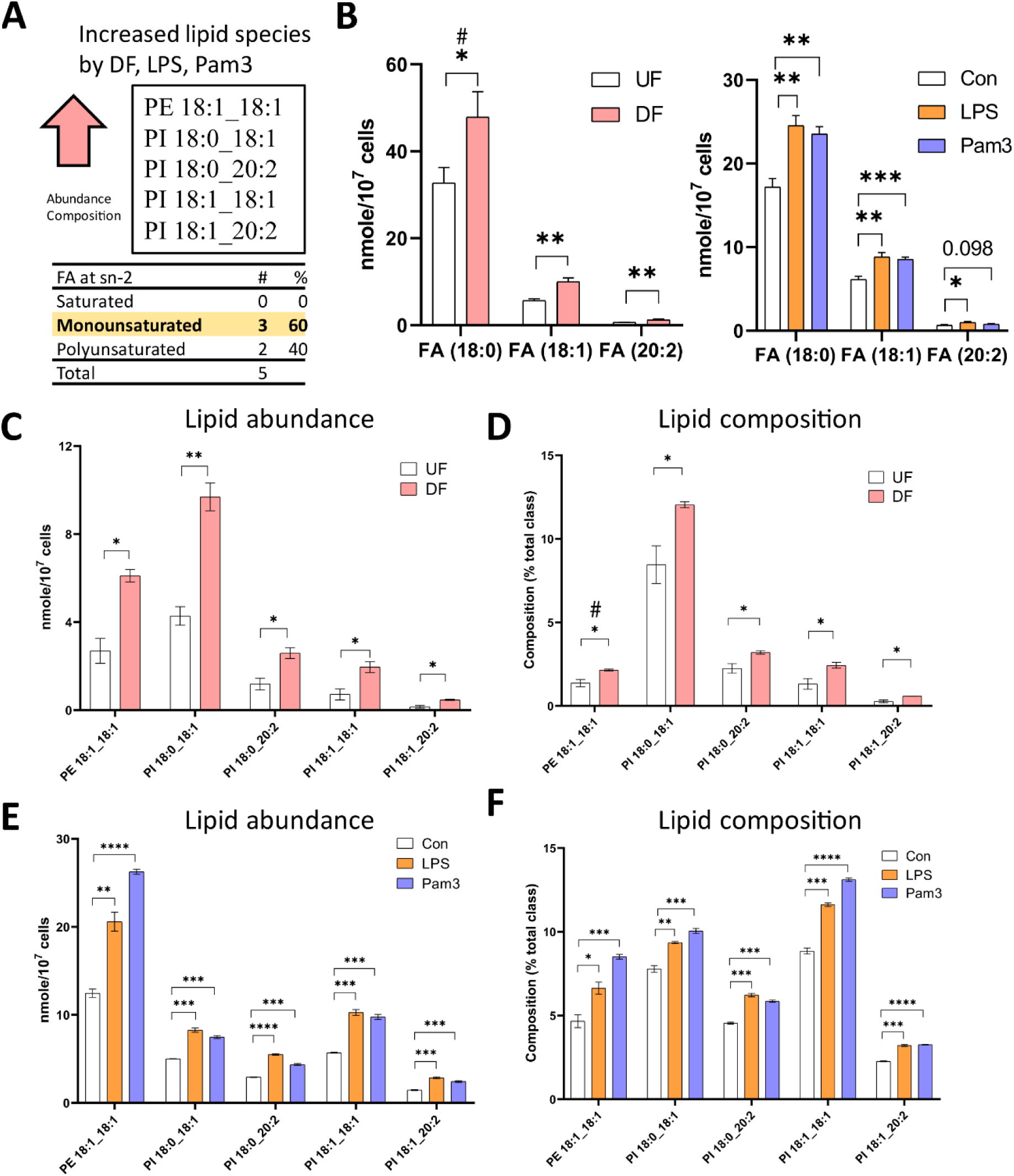
Specific lipid species are increased in HAECs exposed to disturbed flow or inflammatory agonists. **(A)** Summary of cross analysis from the lipidomics datasets. Five lipid species (PE 18:1_18:1, PI 18:0_18:1, PI 18:0_20:2, PI 18:1_18:1, PI 18:1_20:2) that were significantly elevated in abundance and percent composition for HAECs exposed to both DF and inflammatory agonist treatment were identified. **(B)** Abundance of fatty acids incorporated into the lipid species identified in (A) in HAECs under UF and DF or HAECs exposed to vehicle (control), LPS, and Pam3 treatments (n=3). **(C)** Lipid abundance of the shared lipid species in HAECs under UF vs. DF (n=3). **(D)** Lipid composition of the shared lipid species in HAECs under UF vs. DF (n=3). **(E)** Lipid abundance of the shared lipid species in HAECs in response to vehicle, LPS, and Pam3 (n=3). **(F)** Lipid composition of the commonly shared lipid species in HAECs in response to vehicle, LPS, and Pam3 (n=3). Bar graph shown as mean ± SD. *p<0.05, **p<0.01, ***p<0.001, ****p<0.001, ns=not significant by two-tailed unpaired *t*-test. #p<0.05 by one-tailed Mann-Whitney U test.

### Monounsaturated fatty acid incorporation into phospholipids at the sn-2 position is enhanced with disturbed flow

We subsequently investigated the acyl tail compositions of fatty acids incorporated at the sn-2 position in phospholipids for cells exposed to DF and UF. Comparing the altered phospholipid species by abundance, we found that ~61% of the phospholipids contained MUFAs at the sn-2 position, while ~24% contained a saturated fatty acid (SFA) and ~14% contained a polyunsaturated fatty acid (PUFA) **(Figure 6A & 6B)**. Over half of the phospholipids that were significantly changed in DF by composition contained a MUFA and 42% contained a PUFA at the sn-2 position **(Figure 6B & 6C)**.

**Figure 6:**
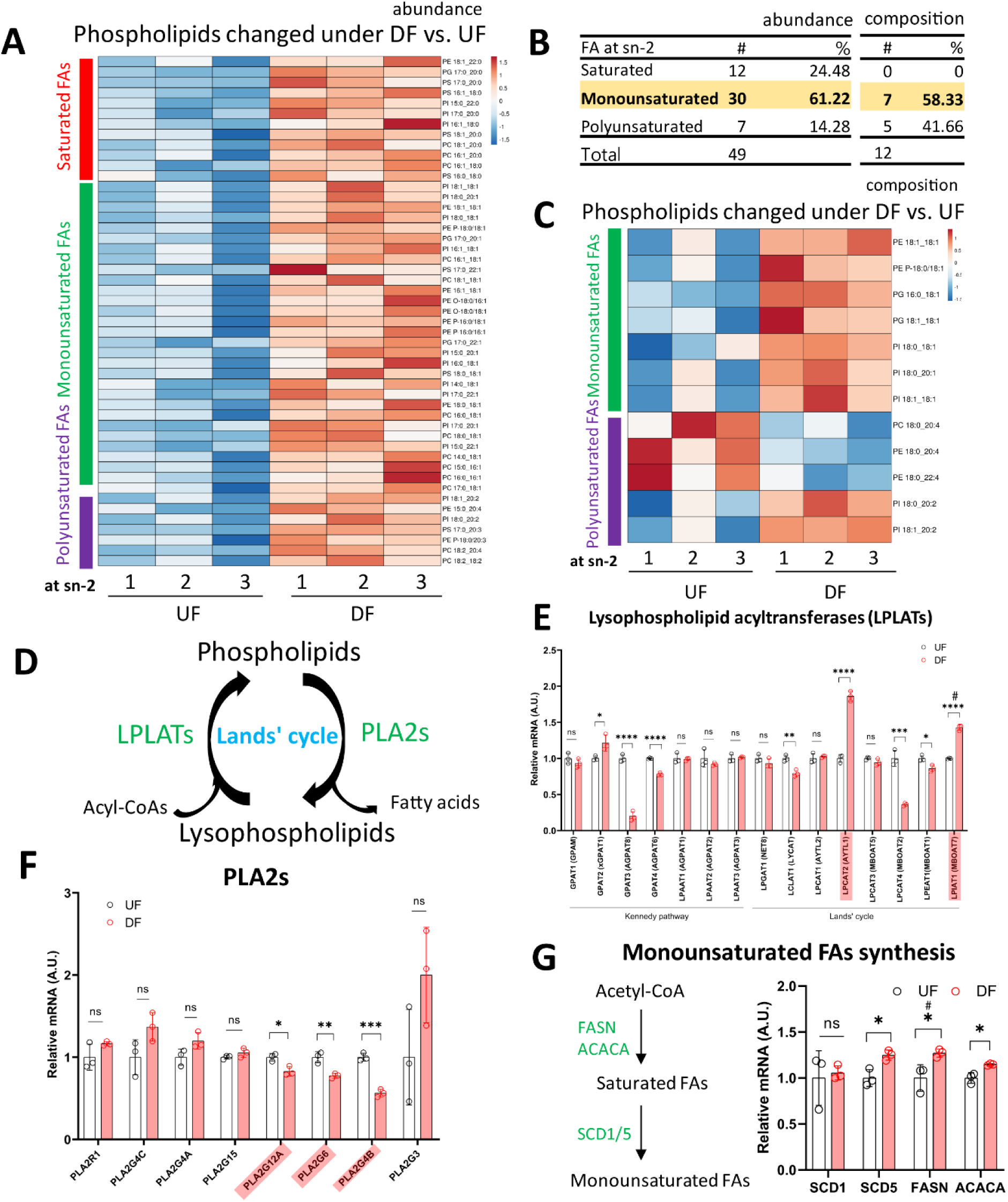
Incorporation of monounsaturated fatty acids into phospholipids at the sn-2 position was enhanced under DF. **(A)** Heatmap of phospholipids altered in abundance under DF vs. UF. **(B)** Table listing the type of fatty acid incorporated at the sn-2 position of altered phospholipids. Values of abundance from (A) and values of composition from (C). **(C)** Heatmap of phospholipids changed in composition under DF vs. UF. (A and C) Color bars represent fatty acid type located at the sn-2 position. Red = saturated, Green = monounsaturated, Purple = polyunsaturated. **(D)** Schematic diagram of Lands’ cycle. Data from RNAseq dataset for HAECs under DF vs. UF showing **(E)** gene expression of Lysophospholipid acyltransferases (LPLATs), **(F)** gene expression of Phospholipase A2 (PLA2s) isoforms and **(G)** genes associated with monounsaturated fatty acid synthesis (n=3). Bar graph shown as mean ± SD. *p<0.05, **p<0.01, ***p<0.001, ****p<0.001, ns=not significant by two-tailed unpaired *t*-test. #p<0.05 by one-tailed Mann-Whitney U test.

Phospholipids are produced by *de novo* synthesis and their fatty acyl chains can subsequently be remodeled by Land’s cycle, which involves removing a fatty acid from the sn-2 position by phospholipase A2 before re-acylating it by lysophospholipid acyltransferases (LPLATs) ^25^ **(Figure 6D)**. Therefore, the observed changes in phospholipid composition of membranes under DF compared to UF could be due to (1) altered uptake or synthesis of fatty acids for *de novo* phospholipid production or (2) changes in phospholipid remodeling.

From the RNASeq data, we observed higher expression of the fatty acid transporter *CD36*, a major route of fatty acid uptake, for HAECs under DF compared to UF (**Figure 2H**). Additionally, the sequencing data indicated that genes involved in SFA and MUFA synthesis, including *FASN*, *ACACA*, and Stearoyl-CoA Desaturase 5 (*SCD5*) were significantly increased for HAECs exposed to DF (**Figure 6G**). The RNASeq data suggest that higher *de novo* MUFA synthesis could contribute to the increased abundance of MUFA-containing phospholipids under DF. The data also showed that the expression levels of select LPLATs were changed under DF compared to UF (**Figure 6E**). Specifically, gene expression of the membrane bound O-acyltransferase domain-containing 7 (*MBOAT7*)/ lysophosphatidylinositol acyltransferase 1 (*LPIAT1*), an enzyme involved in PI remodeling ^25–27^, and lysophosphatidylcholine acyltransferase 2 (*LPCAT2*), which remodels platelet-activating factor (PAF) and PC ^25,26^, were significantly upregulated under DF. Together the observations suggest that changes in fatty acid import and synthesis, combined with changes in the phospholipid remodeling machinery (**Figure 6F**), could contribute to the changes in phospholipid species composition for HAECs exposed to DF compared to UF.

The top 20 phospholipid species used for the PCA loadings for PC1 are listed in **Supplemental Figures 10A & 10B**. Notably, MUFAs were the dominant acyl tails incorporated at the sn-2 position of the top 20 phospholipid species under both DF (65%) and LPS/Pam3 stimulation (45%), as assessed by PCA analysis. Additionally, the abundance of MUFAs in phospholipids was significantly increased after exposure to DF and pro-inflammatory stimuli (**Supplemental Figure 10 C–H**), suggesting a shift towards a program of increased MUFA synthesis and/or FA import.

## DISCUSSION

In this study, we found that the applied flow pattern (DF versus UF) distinctively alters the EC transcriptome and lipidome. While previous reports have shown that laminar shear stress alters the lipidome of pulmonary artery ECs ^28^, this is the first report to correlate how flow pattern affects both metabolic gene expression and lipidomic profiles of arterial ECs. Using RNA sequencing and shotgun lipidomics, we found that HAECs exposed to DF have an inflammatory transcriptional signature and increased cellular lipid content compared to HAECs exposed to UF. Increased lipid accumulation has previously been reported in other cell types exposed to stress conditions, including viral infection-mediated inflammation ^29^. Furthermore, in human microvascular ECs, the expression of the FA transporter CD36 increased in response to viral infection ^30^. Analysis of our RNA sequencing data for DF versus UF revealed that genes involved in fatty acid synthesis (*FASN* and *ACACA*), and fatty acid transport (*CD36* and *FATP1*), were upregulated, while genes involved in lipid and fatty acid catabolic processes were reduced in HAECs. These transcriptional changes suggest that DF may induce changes in the rate of lipid uptake and synthesis to supply more fatty acids for phospholipid and neutral lipid synthesis. Future experiments using stable isotope tracing to measure the contribution of long chain SFA and MUFA synthesis for the observed altered phospholipid composition in response to DF will enable quantification of EC lipid uptake and synthesis.

Exposure of HAECs to DF or an inflammatory agonist, either LPS or Pam3, led to an increase in the abundance of MUFA-containing phospholipids. We noted that genes involved in SFA and MUFA synthesis, including *FASN*, *ACACA* and *SCD5,* were elevated under DF. Stearoyl Coenzyme A (CoA) desaturases incorporates a Δ9-cis-double bond into SFA-CoA to make MUFAs ^31^ and MUFAs have been reported to counteract membrane stress induced by SFAs ^32,33^. Furthermore, MUFAs have been shown to play a role in the modulation of various stress signaling pathways ^31,33–36^. In fact, a recent study in macrophages found that Toll-like receptor-agonist exposure increased *de novo* MUFA synthesis and that attenuating MUFA synthesis with inhibition of SCDs led to heightened inflammation ^37^. Therefore, the observed increase in MUFA-containing phospholipid content in response to DF in ECs could be a similar compensatory mechanism to alleviate the DF-induced inflammatory state. Examination of the mechanisms by which MUFA-containing phospholipid content is increased in response to inflammatory agonists or DF, and the specific pathways that are modulated by MUFA-containing phospholipids, is warranted.

The lipidomics data revealed an increase in the major membrane lipids PC and PE for HAECs exposed to DF compared to UF. Notably, the total unesterified fatty acid content of cells exposed to DF was higher. This increase in PC and PE content could be driven by increased fatty acid import via *CD36* or higher *de novo* fatty acid synthesis, since fatty acid availability drives phospholipid synthesis ^38^. Another possible explanation for the increased pool size of PC and PE is the expansion of cellular organelle volume and/or organelle number in ECs under DF. In fact, athero-susceptible shear stress has been reported to induce ER stress and ER expansion, thereby leading to EC dysfunction and inflammation ^39^. However, future studies are required to investigate these possible scenarios and the functional importance of increased PC and PE content in ECs exposed to DF.

Finally, we identified five lipid species (PE 18:1_18:1, PI 18:0_18:1, PI 18:0_20:2, PI 18:1_18:1, PI 18:1_20:2) that were significantly elevated in abundance and percent composition for HAECs exposed to both DF and inflammatory agonist treatment. We noted that the total abundance of the fatty acids incorporated into those lipid species– stearic acid, oleic acid and eicosadienoic acid– were elevated in cells exposed to DF and exposure to inflammatory agonists. By RNA sequencing, we observed altered expression of genes related to PI signaling and metabolism under DF. While the role of PI metabolism on cellular stress signaling has not been examined in detail in ECs, a recent study showed that the SCD1-derived ‘lipokine’ PI 18:1_18:1 suppressed stress signaling in fibroblasts ^40^. The results in fibroblasts indicated that PI 18:1_18:1 was elevated in response to cytotoxic stressors and prevented p38 MAPK activation and impeded the unfolded protein response ^40^. Future studies are needed to determine how our observed changes in the abundance of PI species in response to DF affects signaling in ECs.

By exposing HAECs to inflammatory agonists, Pam3 or LPS, we confirmed that the EC inflammatory state is associated with higher lipid accumulation. Considering the increased lipid load, it is interesting to consider whether altering lipogenesis in the presence of an inflammatory stimulus would alter the EC phenotype. The previous study in macrophages showed that increasing *de novo* MUFA synthesis modulated the macrophage inflammatory state and that attenuation of MUFA synthesis by inhibition of Stearoyl-CoA Desaturases (SCDs) elevated Toll-like receptor dependent inflammation ^37^. Therefore, it is conceivable that inhibition of *de novo* lipogenesis, a step upstream of SCDs, by pharmacological inhibition or by deletion of Fatty Acid Synthase could also alter the EC phenotype in response to DF. However, we note the critical role of *de novo* lipogenesis in ECs, including its role in the context of angiogenesis, eNOS activity, and permeability ^41,42^. Therefore, inhibition of lipogenesis in the setting of an inflammatory stimulus could have broad off-target effects. Additionally, it is likely that a major contributor to the increase in EC fatty acid content in an inflammatory setting is the increase in CD36, which transports fatty acids into cells and inhibition of lipogenesis alone may not be sufficient to block the increase in cellular fatty acids after exposure to DF. Future studies are needed to dissect the impact of the observed elevation of lipids in response to inflammatory stimuli in ECs.

We note that our in vitro flow system using parallel plate flow chambers does not recapitulate the physiological oscillatory disturbed flow patterns of the aortic arch. However, considering the limitations of in vitro models, we confirmed that using the ibidi u-slide flow chambers under continuous uni-directional laminar flow (20 dynes/cm^2^) and oscillatory flow (4 dynes/cm^2^, 2 Hz) effectively provided athero-protected and athero-prone phenotypes as evidenced by EC cell morphology, gene expression, and protein expression. We also note that HAECs from one single healthy donor were used throughout this study. The choice of a single donor was to provide technical replicates for the changes in gene expression and lipids across treatments. Future work is certainly warranted to investigate how variations in genetics affect the gene expression and lipid responses across multiple donor HAECs in the context of aging, sex, and genetic backgrounds.

In conclusion, we provide transcriptomic and lipidomic datasets of HAECs under athero-protective UF and athero-prone DF patterns. Additionally, we map the lipidome of HAECs after exposure to the inflammatory agonists LPS and Pam3. We anticipate that these datasets will be a useful resource for the vascular biology community to be further dissected and advance our understanding of how flow pattern influences the EC phenotype.

## DATA AVAILABILITY

RNA sequencing data have been deposited in the NCBI’s Gene Expression Omnibus database (GSE266537).

## AUTHOR CONTRIBUTIONS

J.J.M. designed and supervised the study. S.H. and K.J.W performed experiments. S.J.B. shared reagents and experimental expertise. S.H. performed data analysis. S.H., J.P.K. and J.J.M wrote the paper.

## ACKNOWLEDGEMENTS

This work was funded by the National Center for Advancing Translational Sciences UCLA CTSI Grant UL1TR001881 and NIH grant P30 DK063491. S.H. was supported as a Jim Easton CDF Investigator; J.P.K. was supported by AHA Postdoctoral Fellowship 903306; J.J.M was supported by American Heart Association Career Development Award 19CDA34760007.

---

**SUPPLEMENTAL FIGURE 1:**
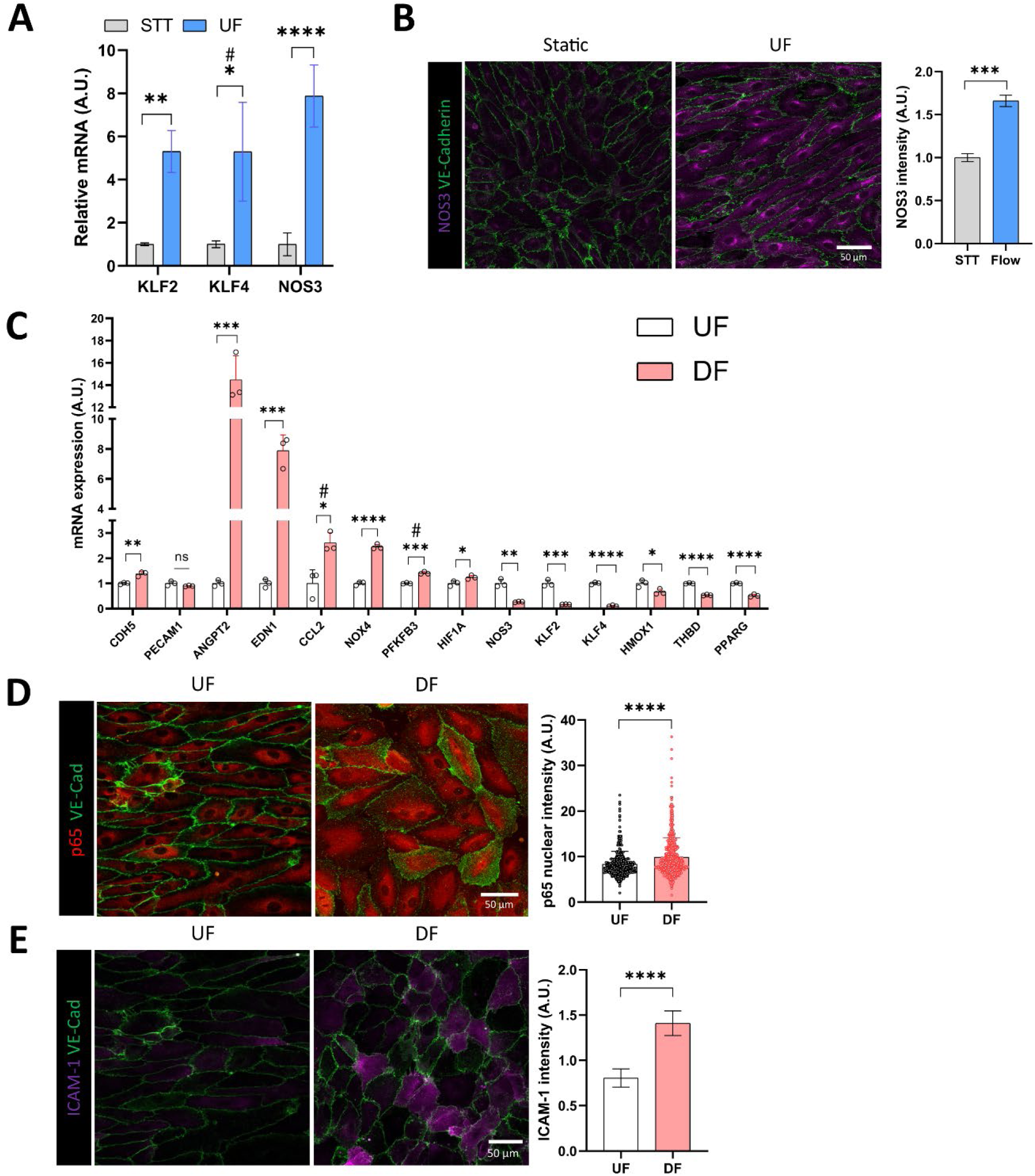
Specific genes altered in response to DF. (A) Expression levels of flow-responsive genes for HAECs exposed to UF or static conditions for 48 h (n=3). (B) Confocal imaging of total eNOS expression in HAECs under static or UF. Elevation of eNOS protein was measured by immunostaining and quantification of fluorescence intensity (n=3). Green = VE-Cadherin, Magenta = eNOS. (C) Expression levels of genes known to change in response to DF vs. UF (n=3). (D) NF-κB p65 nuclear localization, an indicator of enhanced NF-κB-mediated cellular inflammation, was enhanced under DF in HAECs. Red = p65, Green = VE-Cadherin. (E) Representative confocal images of ICAM-1 and VE-Cadherin immunostaining of HAECs exposed to UF vs. DF. ICAM-1 protein expression was greater under DF as quantified by mean fluorescence intensity (n=5). Bar graph shown as mean ± SD. *p<0.05, **p<0.01, ***p<0.001, ****p<0.001, ns=not significant by two-tailed unpaired *t*-test. #p<0.05 by one-tailed Mann-Whitney U test.

**SUPPLEMENTAL FIGURE 2:**
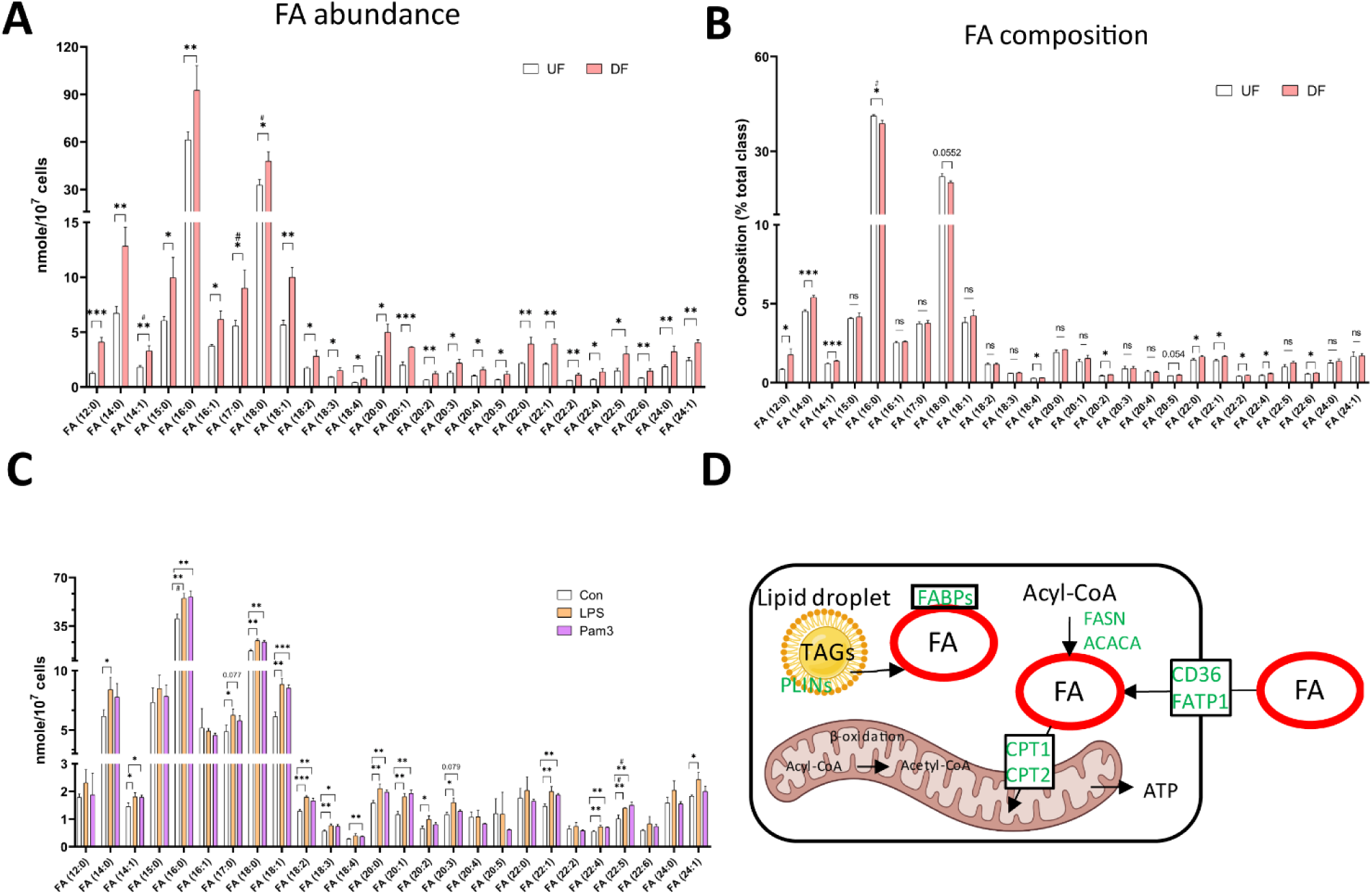
Fatty acid abundance and composition in HAECs under DF versus UF. **(A)** Bar plot of fatty acid abundance in HAECs exposed to UF vs. DF (n=3). **(B)** Bar plot of fatty acid composition in HAECs under UF vs. DF (n=3). **(C)** Bar graph of fatty acid abundance in HAECs exposed to vehicle, LPS, and Pam3. Bar graph shown as mean ± SD (n=3). **(D)** Schematic diagram of the proteins that regulates fatty acid abundance in cells. Fatty acid abundance in cells is regulated by proteins that modulate the uptake, synthesis, and transport of fatty acids, as well as proteins that hydrolyze fatty acids from neutral lipids and phospholipids. *p<0.05, **p<0.01, ***p<0.001, ****p<0.001, ns=not significant by two-tailed unpaired *t*-test. #p<0.05 by one-tailed Mann-Whitney U test.

**SUPPLEMENTAL FIGURE 3:**
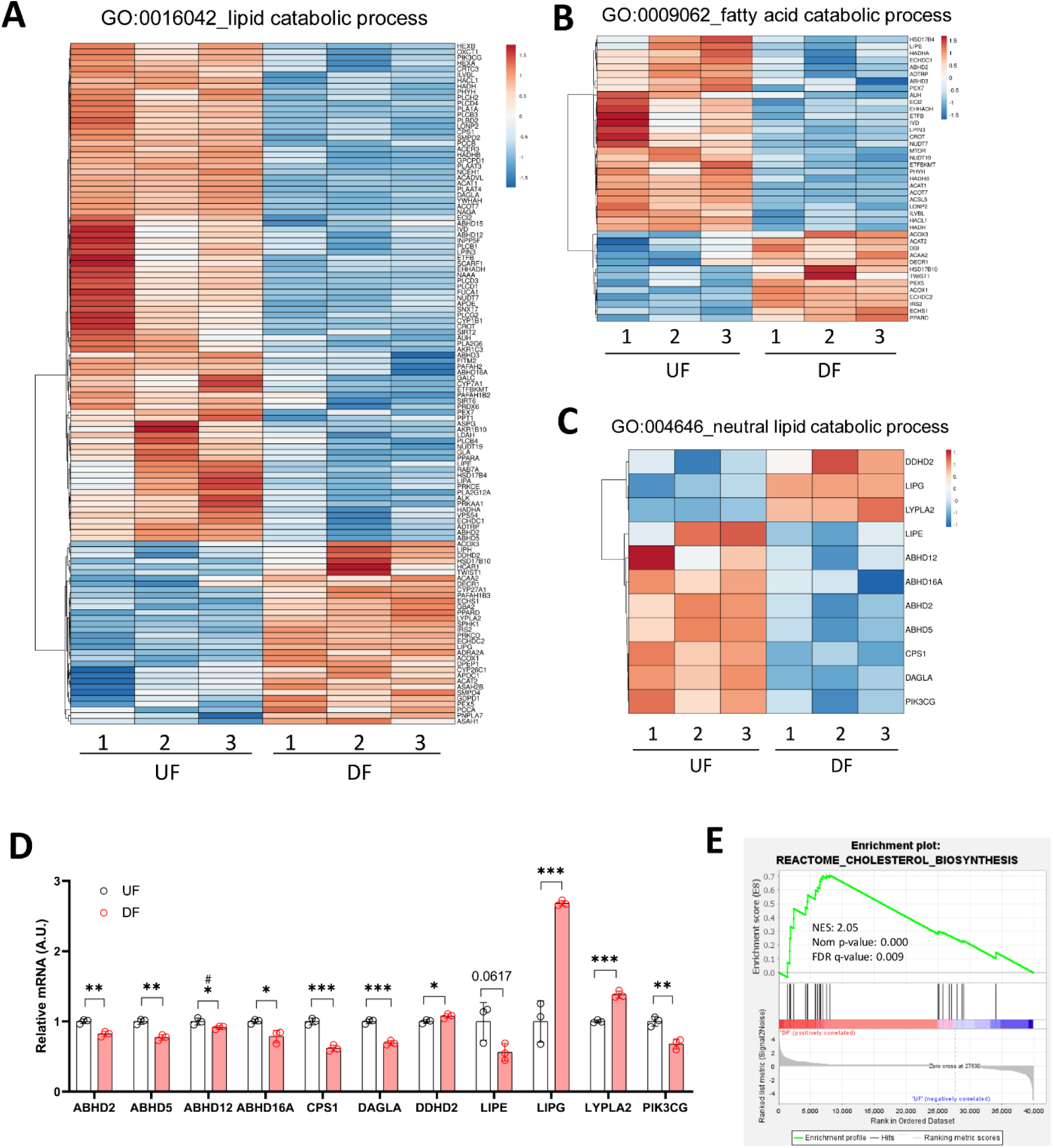
Expression of genes involved in lipid catabolic process. **(A)** Heatmap of genes involved in lipid catabolic process from gene ontology (GO) database (GO:0016042). (**B**) Heatmap of genes involved in fatty acid catabolic process from gene ontology (GO) database (GO:0009062). (**C**) Heatmap of genes involved in neutral lipid catabolic process from gene ontology (GO) database (GO:004646). **(D)** Bar plot using the value from (B) (n=3). (**E**) GSEA enrichment plot using REACTOME database. Genes involved in cholesterol biosynthesis pathway were significantly enriched under DF vs. UF. NES = normalized enrichment score. norm p-value = Normalized p-value. FDR q-value = false discovery rate q-value. Bar graph shown as mean ± SD. *p<0.05, **p<0.01, ***p<0.001, ns=not significant by two-tailed unpaired *t*-test. #p<0.05 by one-tailed Mann-Whitney U test.

**SUPPLEMENTAL FIGURE 4:**
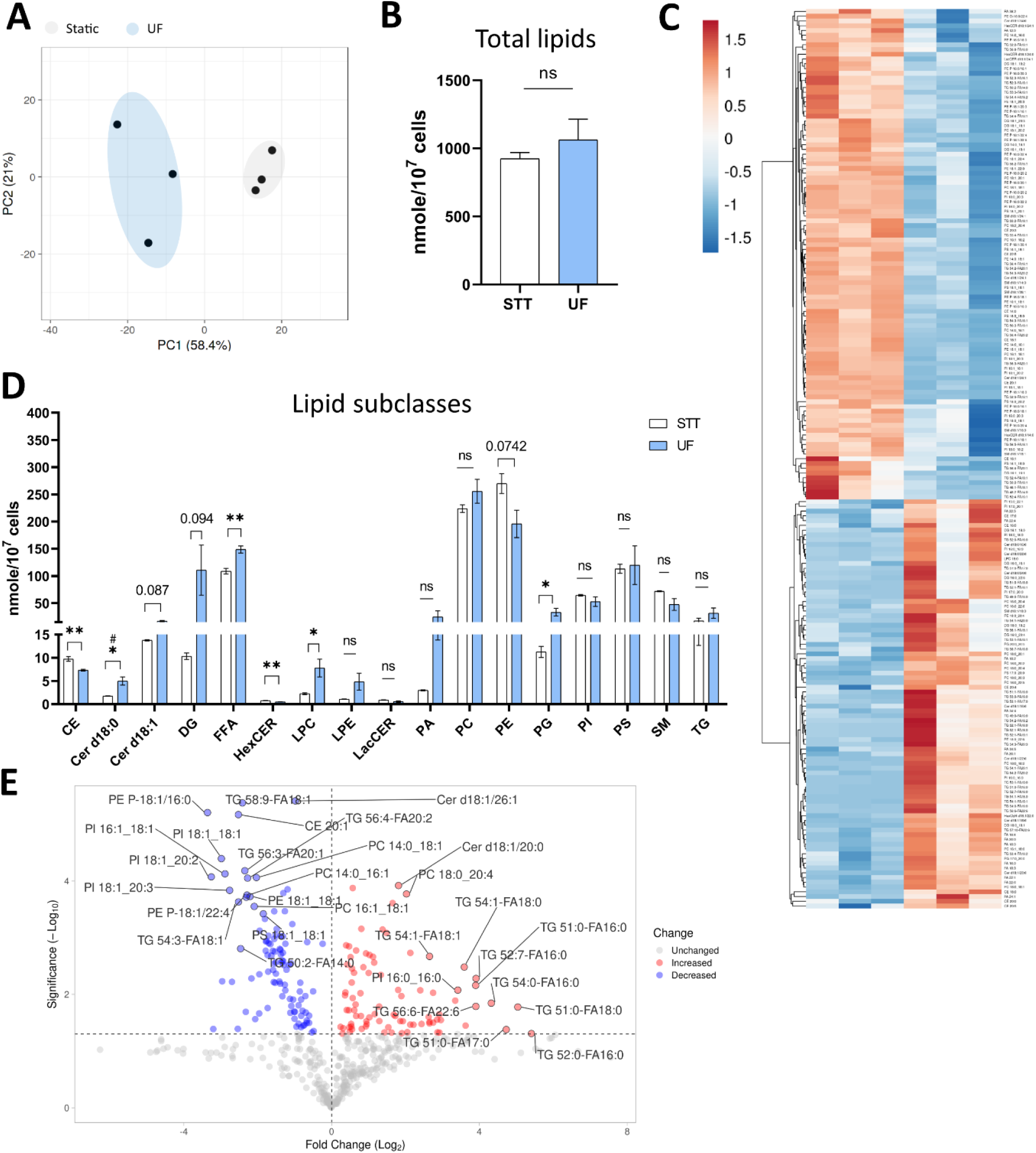
Changes in lipid abundance for HAECs exposed to static versus UF conditions. **(A)** PCA plot of lipid species comparisons for static and UF. Grey = Static control, Blue = UF-exposed HAECs. **(B)** Bar graph for total lipid abundance (n=3). **(C)** Heatmap of altered lipid species under UF vs. static cultured HAECs. **(D)** Bar plot of lipid subclass abundance under static and UF (n=3). **(E)** Volcano plot of lipid species for HAECs exposed to UF vs. static conditions. Bar graph shown as mean ± SD. *p<0.05, **p<0.01, ns=not significant by two-tailed unpaired *t*-test. #p<0.05 by one-tailed Mann-Whitney U test.

**SUPPLEMENTAL FIGURE 5:**
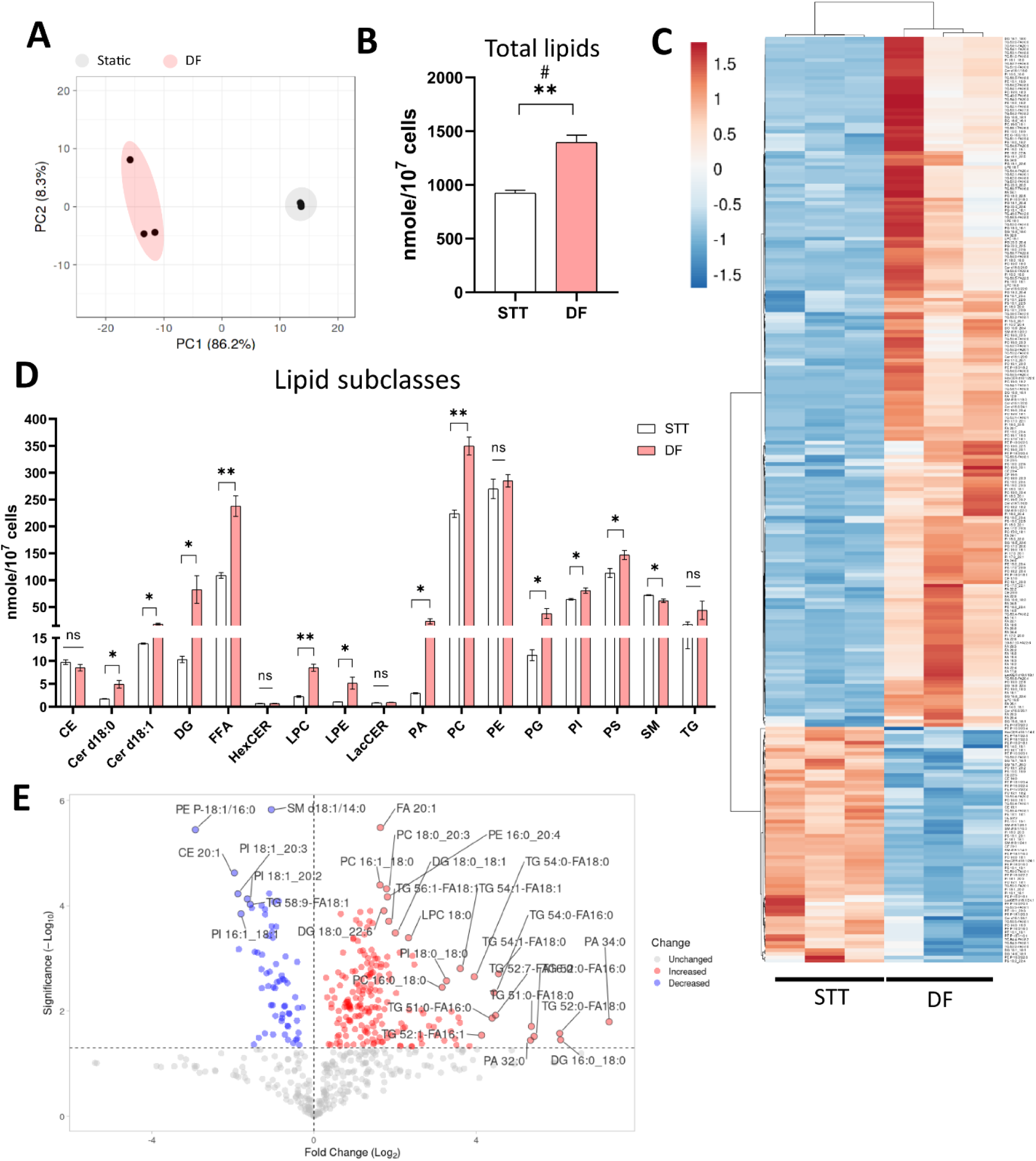
Lipid abundance for HAECs exposed to DF versus static conditions. **(A)** PCA plot using lipid species. Grey = Static control, Red = DF-exposed HAECs. **(B)** Bar graph for total lipid abundance (n=3). **(C)** Heatmap of altered lipid species for HAECs under DF vs. static condition. **(D)** Bar plot of lipid subclass abundance in HAECs exposed to static vs. DF (n=3). **(E)** Volcano plot of lipid species for HAECs under DF vs. static. Bar graph shown as mean ± SD. *p<0.05, **p<0.01, ns=not significant by two-tailed unpaired t-test. #p<0.05 by one-tailed Mann-Whitney U test.

**SUPPLEMENTAL FIGURE 6:**
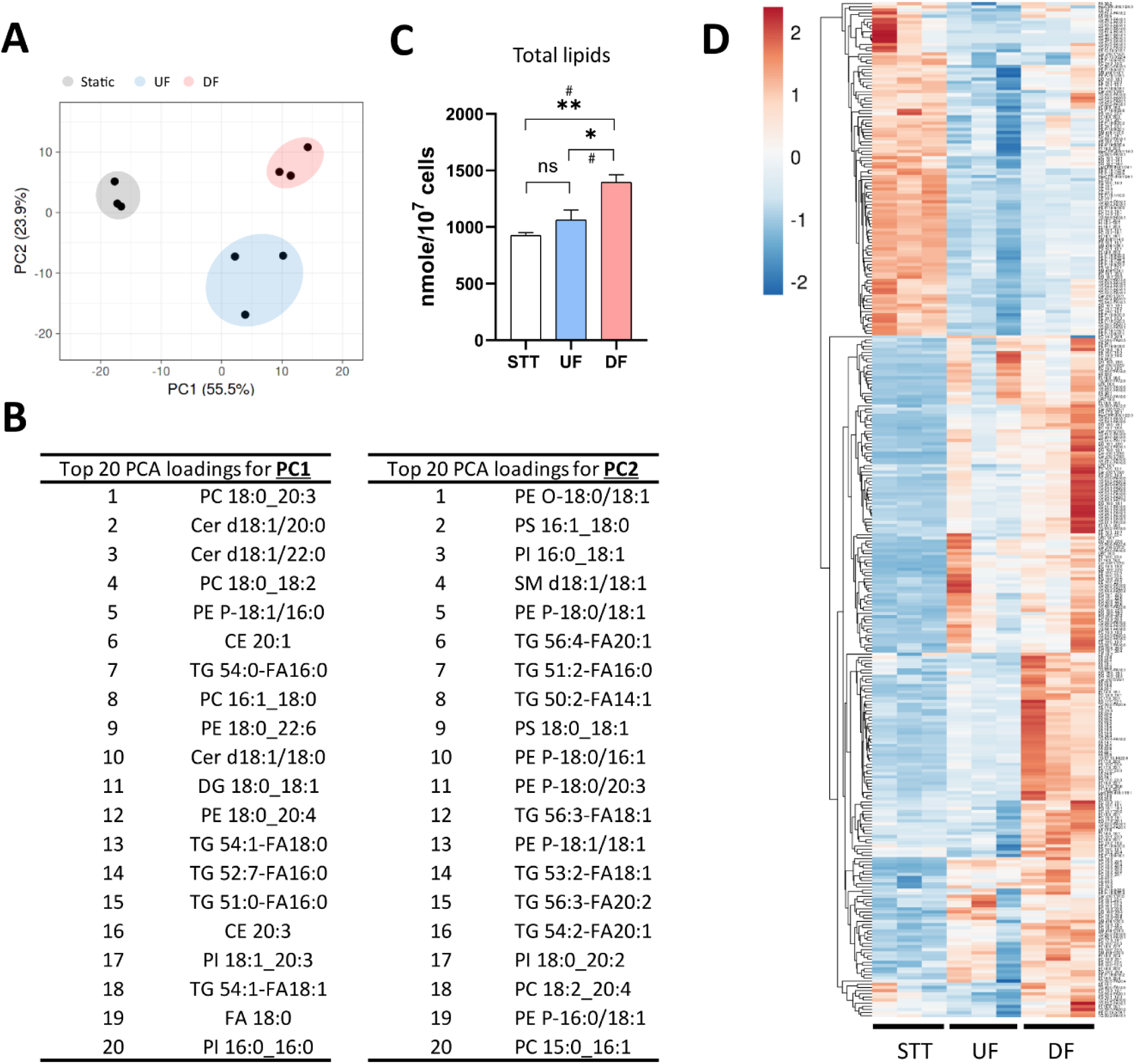
Lipid abundance for HAECs exposed to flow (UF & DF) versus static conditions. **(A)** PCA plot using lipid species. Grey = Static control, Blue = UF-exposed HAECs, Red = DF-exposed HAECs. **(B)** Top 20 PCA loadings for PC1 and PC2. **(C)** Bar graph for total lipid abundance (n=3). **(D)** Heatmap of altered lipid species for HAECs exposed under static vs. UF vs. DF conditions. Bar graph shown as mean ± SD. *p<0.05, **p<0.01, ns=not significant by one-way ANOVA. #p<0.05 by one-tailed Mann-Whitney U test.

**SUPPLEMENTAL FIGURE 7:**
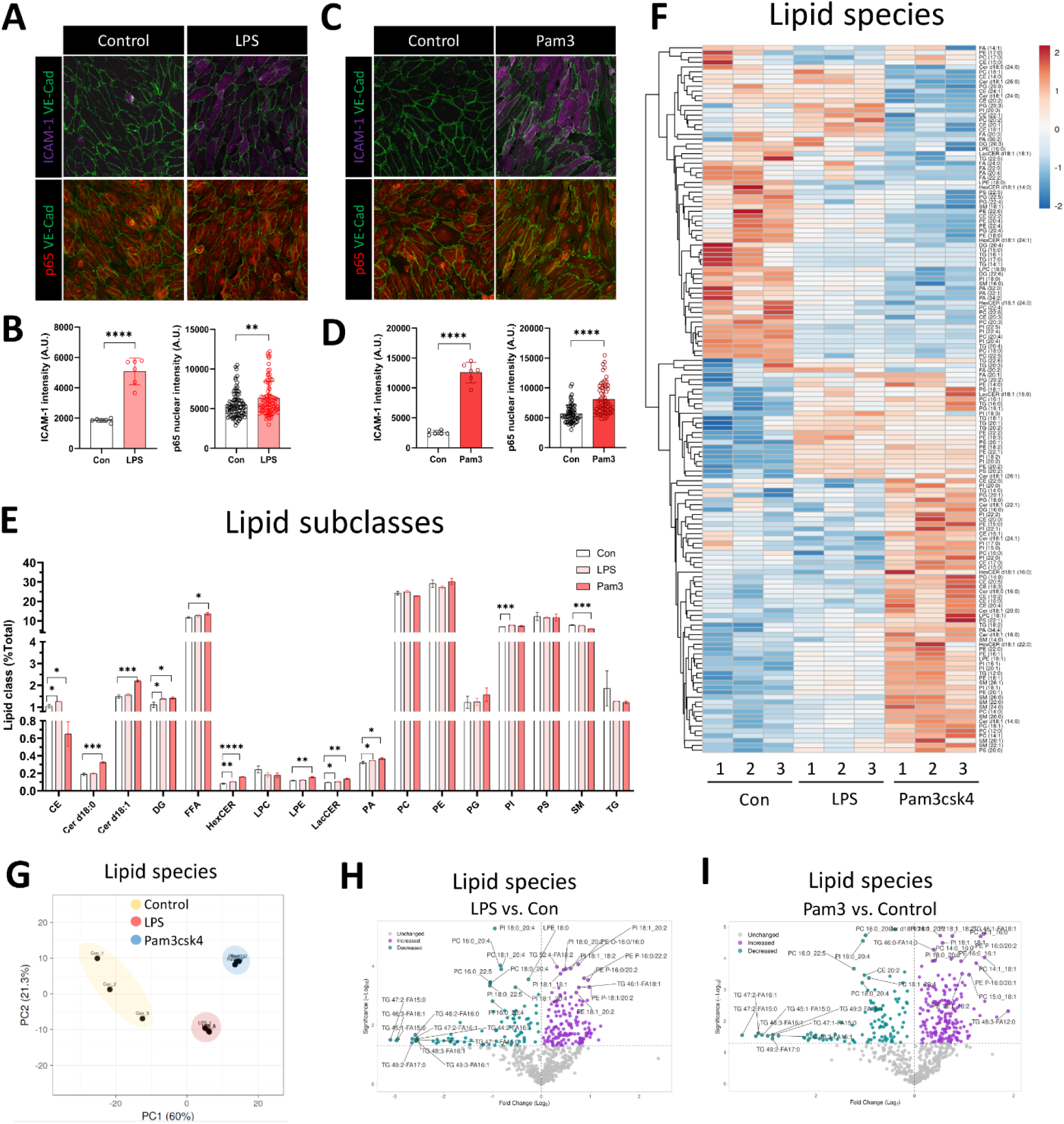
Differential lipid composition in response to inflammatory agonist exposure. **(A)** Confocal imaging of ICAM-1 expression and NF-κB p65 nuclear localization in HAECs in response to control vs. LPS treatment (n=6). Elevation of ICAM-1 expression and p65 nuclear localization were shown in HAECs stimulated with LPS vs. control. Green = VE-Cadherin, Magenta = ICAM-1, Red = p65. **(B)** Dot plots of ICAM-1 and p65 nuclear intensity in control vs. LPS-treated HAECs. **(C)** Confocal imaging of ICAM-1 and NF-κB p65 nuclear localization in HAECs in response to control vs. Pam3 treatment (n=6). Increased ICAM-1 protein expression and p65 nuclear localization were shown in HAECs stimulated with Pam3 vs. control. Green = VE-Cadherin, Magenta = ICAM-1, Red = p65. **(D)** Dot plots of ICAM-1 and p65 nuclear intensity in control vs. Pam3-treated HAECs. (**E**) Bar plot of lipid subclass composition (subclass / total lipid) (n=3). **(F)** Heatmap of altered lipid species composition in HAECs exposed to vehicle (control), LPS, and Pam3. **(G)** PCA plot using lipid species composition dataset. **(H)** Volcano plot of lipid species altered in composition under LPS vs. control. **(I)** Volcano plot of lipid species changed in composition under Pam3 vs. control. Bar graph shown as mean ± SD. *p<0.05, **p<0.01, ***p<0.001, ****p<0.001, ns=not significant by two-tailed unpaired *t*-test.

**SUPPLEMENTAL FIGURE 8:**
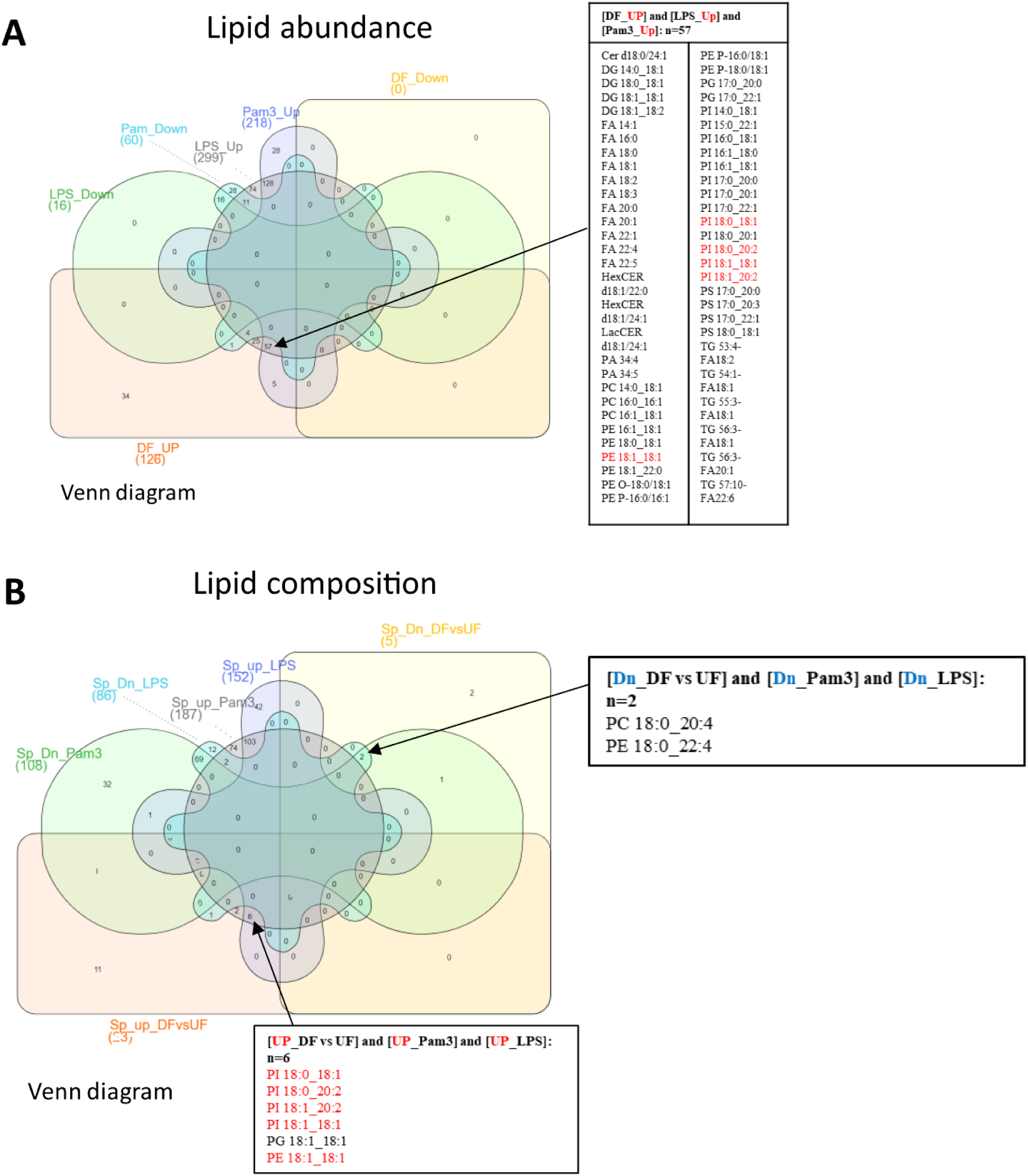
Specific lipid species changed in response to DF and inflammatory agonist exposure. **(A)** Cross analysis using lipid abundance dataset via InteractiVenn. (**B**) Cross analysis using lipid composition dataset via InteractiVenn.

**SUPPLEMENTAL FIGURE 9:**
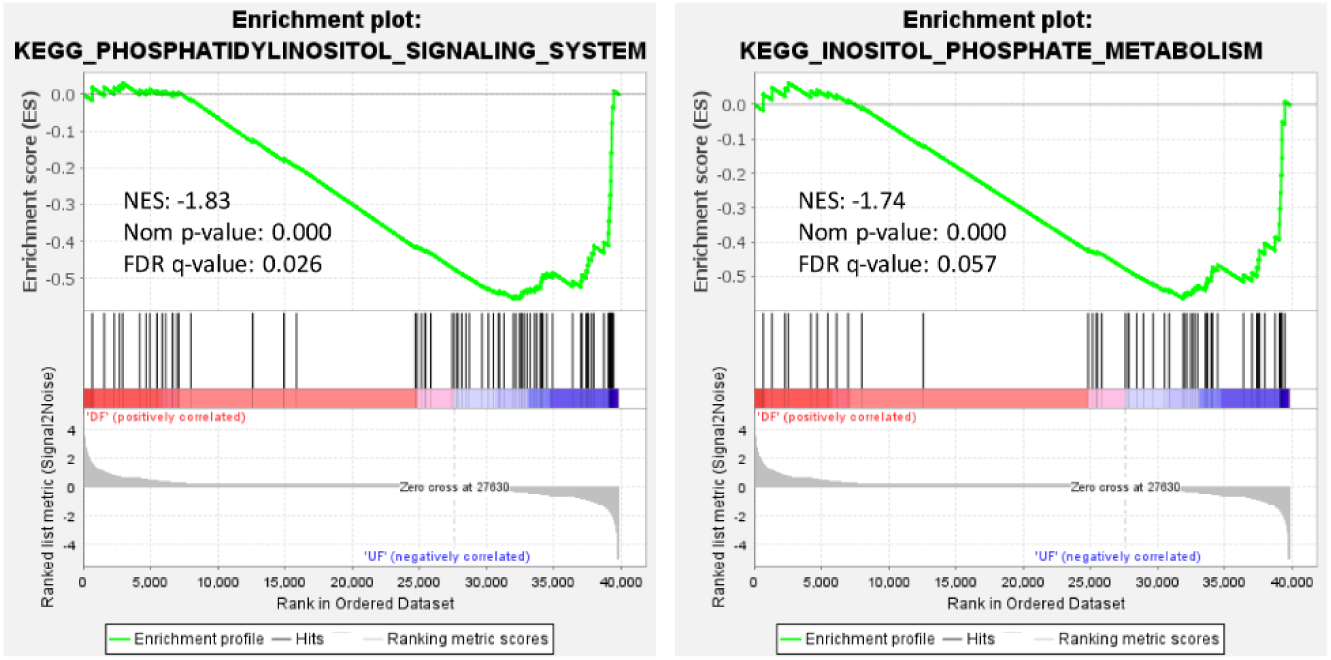
Phosphatidylinositol signaling and metabolism pathways altered under DF. GSEA enrichment plots using KEGG database. Genes involved in phosphatidylinositol (PI) signaling and inositol phosphate metabolism were reduced for HAECs under DF vs. UF. NES = normalized enrichment score. norm p-value = Normalized p-value. FDR q-value = false discovery rate q-value.

**Supplemental Figure 10:**
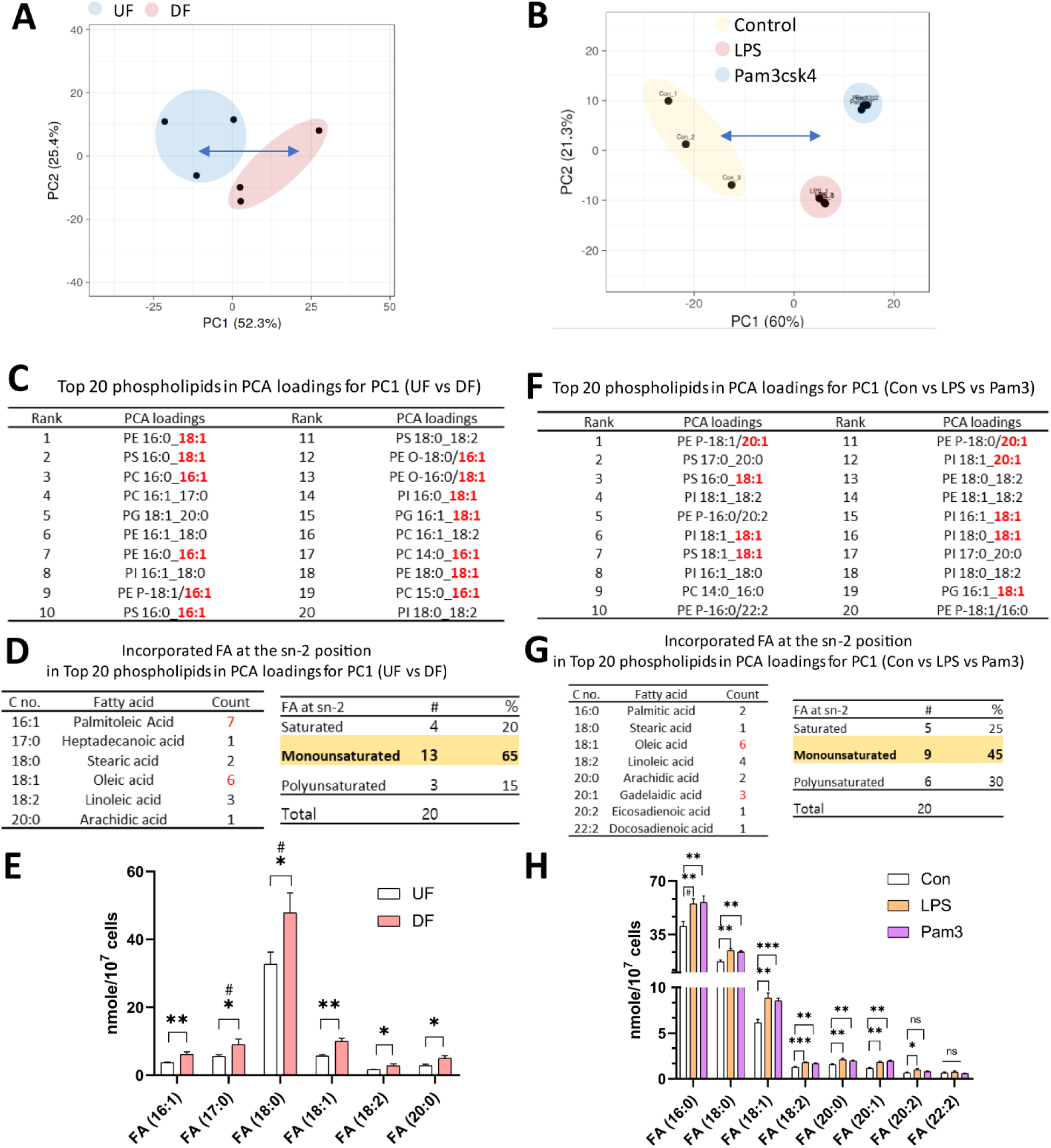
Monounsaturated fatty acids were frequently incorporated into phospholipids at the sn-2 position for HAECs exposed to DF. (**A**) PCA plot of lipid species (by abundance) for HAECs exposed to UF vs. DF. **(B)** PCA plot of lipid species (by abundance) for HAECs under vehicle (control), LPS, and Pam3 treatment. **(C)** Table listing the top 20 phospholipids for PC1 of PCA plot in (A). **(D)** Tables listing the incorporated fatty acid at the sn-2 position and percentage of each type of fatty acid in top 20 phospholipids for PC1 (PCA of UF vs. DF). **(E)** Abundance of fatty acids identified in (D) (n=3). **(F)** Table of the top 20 phospholipids for PC1 (PCA of vehicle vs. LPS vs. Pam3). **(G)** Tables for the number of incorporated fatty acid at the sn-2 position and percentage of each type of fatty acid in the top 20 phospholipids for PC1 (Control vs. LPS vs. Pam3). **(H)** Abundance of fatty acids identified in (G) (n=3). Bar graph shown as mean ± SD. *p<0.05, **p<0.01, ***p<0.001, ns=not significant by two-tailed unpaired *t*-test. #p<0.05 by one-tailed Mann-Whitney U test.

